# Mechanisms involved in follistatin-induced hypertrophy and increased insulin action in skeletal muscle

**DOI:** 10.1101/568097

**Authors:** X. Han, L. L. V. Møller, Estelle De Groote, K.N. Bojsen-Møller, J. Davey, C. Henríquez-Olguin, Z. Li, J.R. Knudsen, T. E. Jensen, S. Madsbad, P. Gregorevic, E. A. Richter, L. Sylow

**Affiliations:** Section of Molecular Physiology, Dept. of Nutrition, Exercise, and Sports, Faculty of Science, University of Copenhagen, Denmark; Faculty of Motor Science, Institute of Neuroscience, Université Catholique Louvain, Belgium; Department of Endocrinology, Copenhagen University Hospital Hvidovre, Hvidovre, Denmark; Center for Muscle Research, Dept. of Physiology, University of Melbourne, Australia

**Keywords:** Muscle wasting, follistatin, TGF-β, glucose uptake, insulin resistance, glycemic control

## Abstract

**Background:** Skeletal muscle wasting is often associated with insulin resistance. A major regulator of muscle mass is the transforming growth factor β (TGF-β) superfamily, including activin A, which causes atrophy. TGF-β superfamily ligands also negatively regulate insulin-sensitive proteins, but whether this pathway contributes to insulin action remains to be determined.

**Methods:** To elucidate if TGF-β superfamily ligands regulate insulin action we used an adeno-associated virus gene editing approach to overexpress the activin A inhibitor, follistatin (Fst288) in mouse muscle of lean and diet-induced obese mice. We determined basal and insulin-stimulated 2 deoxy-glucose uptake using isotopic tracers *in vivo*. Furthermore, to evaluate whether circulating Fst and activin A concentrations are associated with obesity, insulin resistance, and weight loss in humans we analysed serum from morbidly obese subjects before, 1 week, and 1 year after Roux-en-Y gastric bypass (RYGB).

**Results:** Fst288 muscle overexpression markedly increased *in vivo* insulin-stimulated (but not basal) glucose uptake (+75%, p<0.05) and increased protein expression and intracellular insulin signalling of AKT, TBC1D4, PAK1, PDH-E1α, and p70S6K (p<0.05). No correlation was observed between the Fst288-driven hypertrophy and the increase in insulin-stimulated glucose uptake but Fst288 increased basal and insulin-stimulated protein synthesis. Importantly, Fst288 completely normalized muscle glucose uptake in insulin-resistant diet-induced obese mice. RYGB surgery doubled circulating Fst and reduced Activin A (−24%, p<0.05) concentration 1 week after surgery before any significant weight loss in morbidly obese normoglycemic patients, while major weight loss after 1 year did not further change the concentrations.

**Conclusions:** We here present evidence that Fst is a potent regulator of insulin action in muscle and in addition to AKT and p70S6K, we identify TBC1D1, TBC1D4 and PAK1 as Fst targets. A possible role for Fst in regulating glycemic control is suggested because circulating Fst more than doubled post RYGB surgery, a treatment that markedly improved insulin sensitivity. These findings demonstrate the therapeutic potential of inhibiting TGF-β superfamily ligands to improve insulin action and Fst’s relevance to muscle wasting associated insulin resistant conditions in mice and humans.

## Introduction

Skeletal muscle is essential for maintaining an independent lifestyle and muscle strength is inversely associated with all-cause mortality ^1^. Furthermore, skeletal muscle plays a critical role in glycemic control as it accounts up to 75% of insulin-stimulated glucose disposal in humans ^2^. Several common conditions are associated with muscle wasting and/or insulin resistance, including cancer cachexia ^3,4^, sarcopenia ^5,6^, and type 2 diabetes ^2,7,8^. However, there are currently no effective therapeutic strategies to improve muscle wasting and function in such conditions. Restoration of muscle mass, strength, and insulin action could be highly effective in improving these debilitating conditions.

A major regulator of muscle mass is the transforming growth factor β (TGF-β) superfamily, including activin A and myostatin, among which activin A is most relevant to humans ^9^. Activin A and myostatin bind to the activin type II receptor (ActRII) causing atrophy. Clinical trials show that blockade of ActRII using an ActRII antibody (bimagrumab) effectively prevents muscle wasting in sarcopenia ^10^, with marked improvements in insulin sensitivity and HbA1c in insulin resistant individuals ^11^, suggesting that TGF-β signalling integrates muscle mass regulation and insulin action.

Follistatin (Fst) is an endogenously expressed inhibitor of activin A and myostatin. Fst binds to and thereby relieves activin A- or myostatin-induced Smad3-mediated suppression of AKT/mTOR signaling ^12^ and thereby promotes skeletal muscle hypertrophy ^13–15^. Accordingly, Fst has emerged as a potential therapeutic to ameliorate the deleterious effects of muscle wasting ^16–18^ but it remains to be tested whether the action of Fst involves increased muscular insulin action.

Circumstantial evidence suggests that, in addition to treating muscle wasting ^16–18^, targeting Fst could indeed be a novel approach to improve insulin sensitivity in muscle. Recent proteomic profiling has shown that Fst regulates PAK1 ^19^ that is a downstream target of Rac1. Rac1 regulates insulin-stimulated translocation of the glucose transporter, GLUT4 ^20,21^ and this pathway is compromised in muscles from obese and type 2 diabetic patients ^22^. Furthermore, Fst increases signaling through the AKT/mTOR pathway in unstimulated mouse muscle ^12^, likely due to upregulation of Akt and mTOR protein expression ^17^. Thus, Fst affects insulin regulated proteins but it is unknown if this translates into increased insulin-stimulated intracellular signaling and glucose uptake and whether Fst could be a target to ameliorate muscle wasting-associated insulin resistance.

Inspired by these findings, we hypothesized that Fst could be a central regulator of muscular insulin sensitivity in addition to its known effects on muscle mass. We confirmed this hypothesis by showing that Fst overexpression markedly increased insulin sensitivity and completely normalized glucose uptake in insulin resistant mouse muscle. Furthermore, Roux-en-Y gastric bypass (RYGB), a treatment that rapidly and markedly improves insulin sensitivity and glycemic regulation, increased circulating Fst together with a decrease in the ActRII ligand activin A. These findings demonstrate the therapeutic potential of Fst and of inhibiting TGF-β superfamily ligands and highlight the relevance of this pathway to insulin resistance in humans.

## Methods

### Animals

Female C57BL/6J (Taconic, Lille Skensved, Denmark) mice age 20-22 weeks were maintained on a 12 h:12 h light–dark cycle, and group housed at 21-23°C with nesting materials. A 10-week diet intervention was started at 12 weeks of age and mice received either a standard rodent chow diet (Altromin no. 1324; Brogaarden, Horsholm, Denmark), or a 60E% high-fat diet (HFD; Research Diets no. D12492; Brogaarden, Denmark) and water ad libitum. Body weight was assessed biweekly. All experiments were approved by the Danish Animal Experimental Inspectorate. The animal experiments followed the European convention for protection of vertebrate animals used for experiments and other scientific purposes.

### Production of rAAV6 vectors

Generation of cDNA constructs encoding the short isoform of human follistatin, known as Fst288 and subcloning into an adeno-associated (AAV) expression plasmid consisting of a CMV promoter/enhancer and SV40 poly-A region flanked by AAV2 terminal repeats (pAAV2), has been described previously ^12^. Co-transfection of HEK293T cells with the above mentioned pAAV2 and the pDGM6 packaging plasmid generated type-6 pseudotyped viral vectors that were harvested and purified as described previously ^23^ and are herein referred to as rAAV6:Fst288. As a control, a non-coding vector was also generated (rAAV6:MCS). In short, HEK293T cells were plated at a density of 3.2–3.8 × 10^6^ cells onto a 15-cm culture dish 8–16 h before transfection with 10 μg of a vector-genome-containing plasmid and 20 μg of the packaging/helper plasmid pDGM6, by means of the calcium phosphate precipitate method to generate pseudotype 6 vectors. 72 h after transfection, the media and cells were collected and homogenized through a microfluidizer (Microfluidics) before 0.22 μm clarification (Millipore). The vector was purified from the clarified lysate by affinity chromatography over a HiTrap heparin column (GE Healthcare), and ultracentrifuged overnight before resuspension in sterile physiological Ringer’s solution. The purified vector preparations were titered with a customized sequence-specific quantitative PCR-based reaction (Applied Biosystems) ^12^.

### Transfection of mouse muscles using rAAV6 vectors

Viral particles were diluted in Gelofusine (B. Braun, Germany) (Tibialis anterior: 30 μL, Gastrocnemius: 50 μL) to a dosage of 1×10^9^ vector genomes for both muscles. For muscle specific delivery of rAAV6 vectors, 18-20-weeks old mice were placed under general anesthesia (2% isofluorane in O2) and mice were injected intramuscularly with rAAV6:Fst288 in the muscles of one leg and control rAAV6:MCS (empty vector) in the contralateral leg. Mice were terminated 2 weeks after rAAV6 administration.

### Body composition analysis

For the HFD study, fat mass and lean body mass changes were determined by quantitative magnetic resonance using an Echo MRI scanner (EchoMRI-4in1TM, Echo Medical System LLC, Texas, USA) at baseline and week 9 into the diet intervention.

### In vivo 2 deoxy-glucose experiments

To determine 2 deoxy-glucose (2DG) uptake in muscle, [^3^H]2DG (Perkin Elmer) was injected retro-orbitally in a bolus of saline containing 66.7 μCi mL^−1^ [^3^H]2DG (6 μL g^−1^ body weight) in chow and HFD fed mice. The injectate also contained 0.3 U kg^−1^ body weight insulin (Actrapid; Novo Nordisk, Bagsværd, Denmark) or a comparable volume of saline. Prior to stimulation, mice were fasted for 3-5 h from 07:00 h and anaesthetized (intraperitoneal injection of 7.5/9 mg (chow/HFD) pentobarbital sodium 100 g^−1^ body weight 15 minutes before the retro-orbital injection). Blood samples were collected from the tail vein immediately prior to insulin or saline injection and after 5 and 10 min and analyzed for glucose concentration using a glucometer (Bayer Contour; Bayer, Münchenbuchsee, Switzerland). After 10 min, tibialis anterior and gastrocnemius muscles were excised and quickly frozen in liquid nitrogen and stored at −80°C until processing. Once tissues were removed, blood was collected by punctuation of the heart, centrifuged and plasma frozen at −80°C. Plasma samples were analyzed for insulin concentration and specific [^3^H]2DG tracer activity. Plasma insulin was analyzed in duplicate (Mouse Ultrasensitive Insulin ELISA, #80-INSTRU-E10, ALPCO Diagnostics, USA). Tissue specific 2DG uptake was analyzed as described in ^24–26^.

### Glycogen content

Muscle glycogen content in tibialis anterior muscle was determined as glycosyl units after acid hydrolysis as previously described ^27^.

### Cell cultures and estimation of protein synthesis in vitro

L6 muscle cells stably over-expressing c-myc epitope-tagged GLUT4 (L6-GLUT4myc), a kind gift from Amira Klip, were grown in α-MEM medium (Gibco #22571-020) with 10% fetal bovine serum (Sigma-Aldrich #F0804), 100 units mL^−1^ penicillin, 100 μg mL^−1^, and 0.25 μg mL^−1^ Gibco Amphotericin B (Gibco # 15240-062) (5% CO_2_, 37°C). Cells were seeded in 12-well plates and differentiated into multinucleated myotubes in α-MEM media with 2% fetal bovine serum, 100 units mL^−1^ penicillin, 100 μg mL^−1^, and 0.25 μg mL^−1^ Gibco Amphotericin B. AAV-containing medium was added to day 2 myotubes and consisted of 5×10^10^ vector genomes of rAAV6:Fst288 per mL differentiation medium or rAAV6:MCS as a control. Protein synthesis in L6-GLUT4myc cells was analyzed in day 7 myotubes and measured using the non-isotopic SUnSET (SUrface SEnsing of Translation) technique ^28^. For estimation of protein synthesis, cells were serum starved for four hours and incubated with 1 μM puromycin (Gibco # A11138-03) with or without 10 nM insulin (submaximal dose; Actrapid; Novo Nordisk, Bagsværd, Denmark) for 60 min. Immediately following stimulation, cells were chilled on ice, washed in PBS (Gibco #14190-094) and lysed in ice-cold RIPA buffer (Sigma-Aldrich #R0278) supplemented with 20 mM β-glycerophosphate, 10 mM NaF, 2 mM phenylmethylsulfonyl fluoride (PMSF), 2 mM Na3VO4, 10 μg mL^−1^ leupeptin, 10 μg mL^−1^ aprotinin, 3 mM benzamidine. Supernatants were collected by centrifugation (9,000 x g) for 10 min at 4°C. Each experiment was assayed in duplicates or quadruplicates and repeated three times. Puromycin levels were determined as described for immunoblotting. *Human serum Fst and activin A analysis*. Fasting blood samples were obtained from 9 obese glucose-tolerant subjects (3 men, 6 women) age 40.1 ±2.8 years, body weight 116.9 ±4.9 kg) scheduled for laparoscopic RYGB at Hvidovre Hospital (Denmark) after a mandatory diet-induced weight loss of minimum 8% (weight loss was 9.2 ± 1.2%) pre-surgery and at 1 week and 1 year post-surgery. Detailed description of the study design and subject characteristics have been reported elsewhere ^29^. For the present study, we used samples from the nine glucose tolerant patients that completed the 1 year study visit. Serum was stored at −80°C and Fst and activin A analyzed in duplicates using DFN00 and DAC00B ELISA kits, respectively from R&D Systems performed according to manufacturer’s instructions. Clinical experiments were approved by the Ethics Committee of Copenhagen and complied with the ethical guidelines of the Declaration of Helsinki II 2000. Written informed consent was obtained from all participants prior to entering the study. The clinical study is registered at www.ClinicalTrials.gov (NCT01202526).

### Muscle tissue processing

Mouse muscles were pulverized in liquid nitrogen and homogenized 2 × 0.5 min at 30 Hz using a TissueLyser II bead mill (Qiagen, USA) in ice-cold homogenization buffer (10% glycerol, 1% NP-40, 20 mM sodium pyrophosphate, 150 mM NaCl, 50 mM HEPES (pH 7.5), 20 mM β-glycerophosphate, 10 mM NaF, 2 mM phenylmethylsulfonyl fluoride (PMSF), 1 mM EDTA (pH 8.0), 1 mM EGTA (pH 8.0), 2 mM Na3VO4, 10 μg mL^−1^ leupeptin, 10 μg mL^−1^ aprotinin, 3 mM benzamidine). After rotation end-over-end for 30 min at 4°C, supernatants were collected by centrifugation (10,854 x g) for 20 min at 4°C.

### Immunoblotting

Lysate protein concentrations were measured using the bicinchoninic acid method with bovine serum albumin (BSA) as standard (Pierce). Total protein and phosphorylation levels of relevant proteins were determined by standard immunoblotting techniques loading equal amounts of protein. The primary antibodies used are presented in Table 1. Protein levels of the PDH-E1α subunit, PDH-E1α phosphorylation at Ser293 and Ser300 and protein levels of PDK4 were determined using antibodies, as previously described ^30,31^, all kindly provided by D. Grahame Hardie. Polyvinylidene difluoride membranes (Immobilon Transfer Membrane; Millipore) were blocked in Tris-buffered saline (TBS)-Tween 20 containing 3% milk or 5% BSA protein for 30–60 min at room temperature. Membranes were incubated with primary antibodies overnight at 4°C, followed by incubation with horseradish peroxidase-conjugated secondary antibody for 30 min at room temperature. Coomassie brilliant blue staining was used as a loading control ^32^. Bands were visualized using the Bio-Rad ChemiDoc MP Imaging System and enhanced chemiluminescence (ECL+; Amersham Biosciences).

**Table 1.**
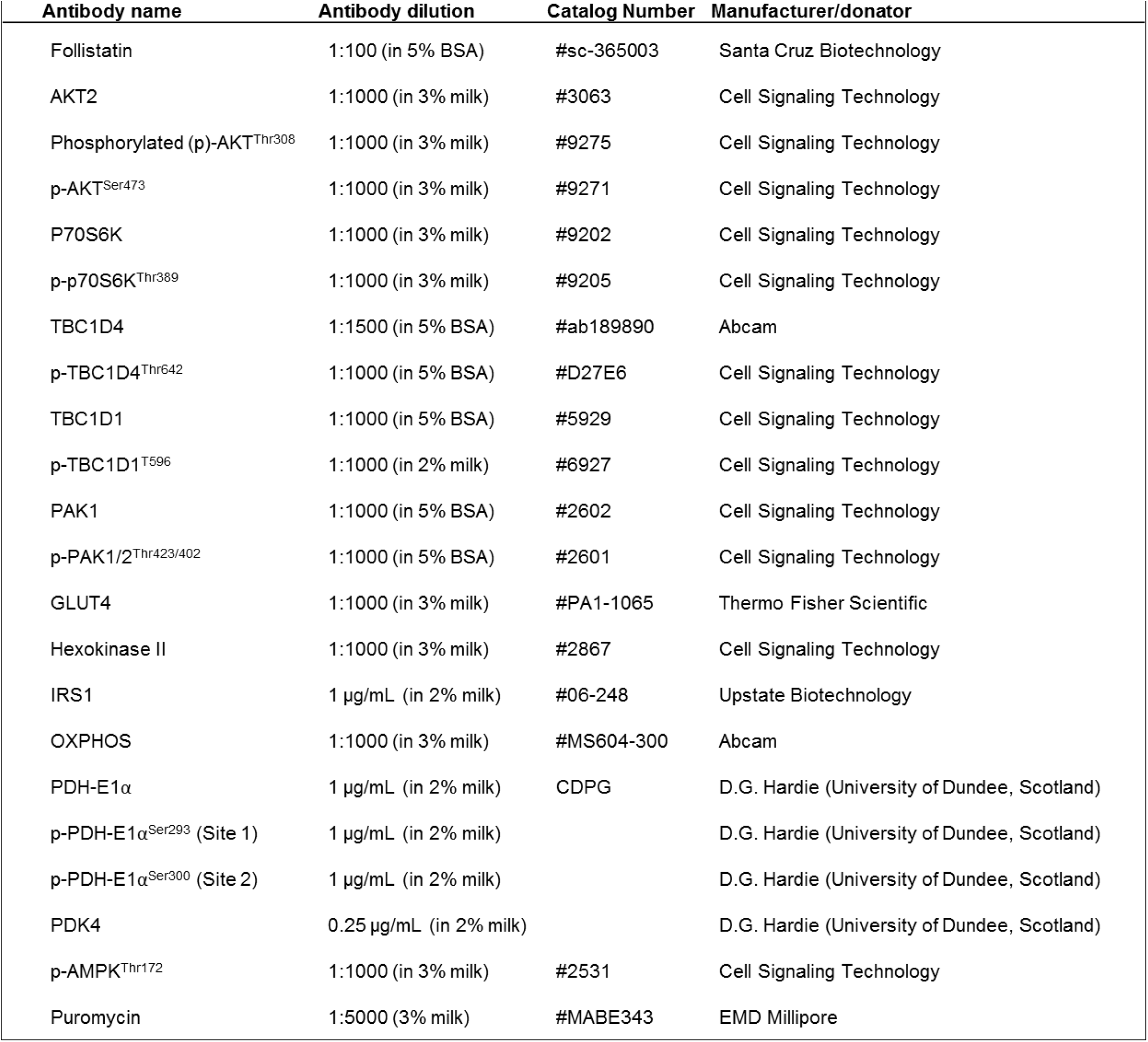
Antibody Table

| **Table 1. Antibody Table** |  |  |  |
| --- | --- | --- | --- |
| **Antibody name** | **Antibody dilution** | **Catalog Number** | **Manufacturer/donor** |
| Follistatin | 1:100 (in 5% BSA) | #sc-365003 | Santa Cruz Biotechnology |
| AKT2 | 1:1000 (in 3% milk) | #3063 | Cell Signaling Technology |
| Phosphorylated (p)-AKT<sup>Thr308</sup> | 1:1000 (in 3% milk) | #9275 | Cell Signaling Technology |
| p-AKT<sup>Ser473</sup> | 1:1000 (in 3% milk) | #9271 | Cell Signaling Technology |
| P70S6K | 1:1000 (in 3% milk) | #9202 | Cell Signaling Technology |
| p-p70S6K<sup>Thr389</sup> | 1:1000 (in 3% milk) | #9205 | Cell Signaling Technology |
| TBC1D4 | 1:1500 (in 5% BSA) | #ab189890 | Abcam |
| p-TBC1D4<sup>Thr642</sup> | 1:1000 (in 5% BSA) | #D27E6 | Cell Signaling Technology |
| TBC1D1 | 1:1000 (in 5% BSA) | #5929 | Cell Signaling Technology |
| p-TBC1D1<sup>T596</sup> | 1:1000 (in 2% milk) | #6927 | Cell Signaling Technology |
| PAK1 | 1:1000 (in 5% BSA) | #2602 | Cell Signaling Technology |
| p-PAK1/2<sup>Thr423/402</sup> | 1:1000 (in 5% BSA) | #2601 | Cell Signaling Technology |
| GLUT4 | 1:1000 (in 3% milk) | #PA1-1065 | Thermo Fisher Scientific |
| Hexokinase II | 1:1000 (in 3% milk) | #2867 | Cell Signaling Technology |
| IRS1 | 1 μg/mL (in 2% milk) | #06-248 | Upstate Biotechnology |
| OXPHOS | 1:1000 (in 3% milk) | #MS604-300 | Abcam |
| PDH-E1α | 1 μg/mL (in 2% milk) | CDPG | D.G. Hardie (University of Dundee, Scotland) |
| p-PDH-E1α<sup>Ser293</sup> (Site 1) | 1 μg/mL (in 2% milk) | | D.G. Hardie (University of Dundee, Scotland) |
| p-PDH-E1α<sup>Ser300</sup> (Site 2) | 1 μg/mL (in 2% milk) | | D.G. Hardie (University of Dundee, Scotland) |
| PDK4 | 0.25 μg/mL (in 2% milk) | | D.G. Hardie (University of Dundee, Scotland) |
| p-AMPK<sup>Thr172</sup> | 1:1000 (in 3% milk) | #2531 | Cell Signaling Technology |
| Puromycin | 1:5000 (3% milk) | #MABE343 | EMD Millipore |

### Statistical analyses

Results are shown as mean ± standard error of the mean (SEM) with the individual values shown when applicable. For figures with a paired design (rAAV6:MCS vs rAAV6:Fst288 treated legs within the same mouse), paired values are connected with a line. Statistical testing was performed using unpaired t-test, paired t-test, and two-way (repeated measures when appropriate) ANOVA as applicable. Sidak post hoc test was performed for multiple comparisons when ANOVA revealed significant main effects or interactions. Pearson’s correlation was used to test the relationship between the change in muscle weight and insulin-stimulated glucose uptake with Fst288 overexpression. Statistical analyses were performed using GraphPad Prism, version 7 (GraphPad Software, La Jolla, CA, USA). The significance level was set at α = 0.05.

## Results

### Insulin-stimulated glucose uptake is increased in Fst288 overexpression-induced hypertrophic muscles

Mice were injected with rAAV6:Fst288 into the muscles of one leg while the contralateral leg was injected with an empty viral vector, rAAV6:MCS (Fig. 1A). Two weeks after the injection, Fst protein expression was markedly increased compared to the control muscle in tibialis anterior (TA) and gastrocnemius muscles injected with rAAV6:Fst288 (Fig. 1B). Fst288 delivery increased TA muscle mass by 33% on average compared to the control muscle (Fig. 1C). This was similar to previous reports ^12^. Mice (one Fst288 treated leg and one control leg) were stimulated with an insulin dose of 0.3 U kg^−1^ as described in ^26^, which lowered blood glucose concentration by 3 mM at the 10 minutes time point compared to saline injected mice (Fig. 1D). Fst288 overexpression resulted in almost a doubling of insulin-stimulated glucose uptake in TA compared to the control muscle (Fig. 1E). Similarly, in gastrocnemius muscle insulin-stimulated glucose uptake was 72% higher in rAAV6:Fst288 treated muscles compared to the contralateral rAAV6:MCS-treated control 2 weeks after rAAV administration (Fig. 1F). Basal (saline injected) glucose uptake was not affected by rAAV6:Fst288. Interestingly, we observed no correlation between the change in muscle mass and insulin-stimulated glucose uptake with Fst288 overexpression in muscle (Fig. 1G), suggesting that Fst288 overexpression per se, rather than the hypertrophy magnitude, drives the increased insulin-stimulated glucose uptake. To investigate the direct effect of Fst288 overexpression on the anabolic response to insulin in muscle, insulin-stimulated puromycin incorporation into L6 myotubes was analyzed. It was found that both basal and insulin-stimulated puromycin incorporation was increased (basal: +40%; insulin: +33%) (Fig. 1H), suggesting that Fst288 led to a general increase in protein synthesis.

**Figure 1.**
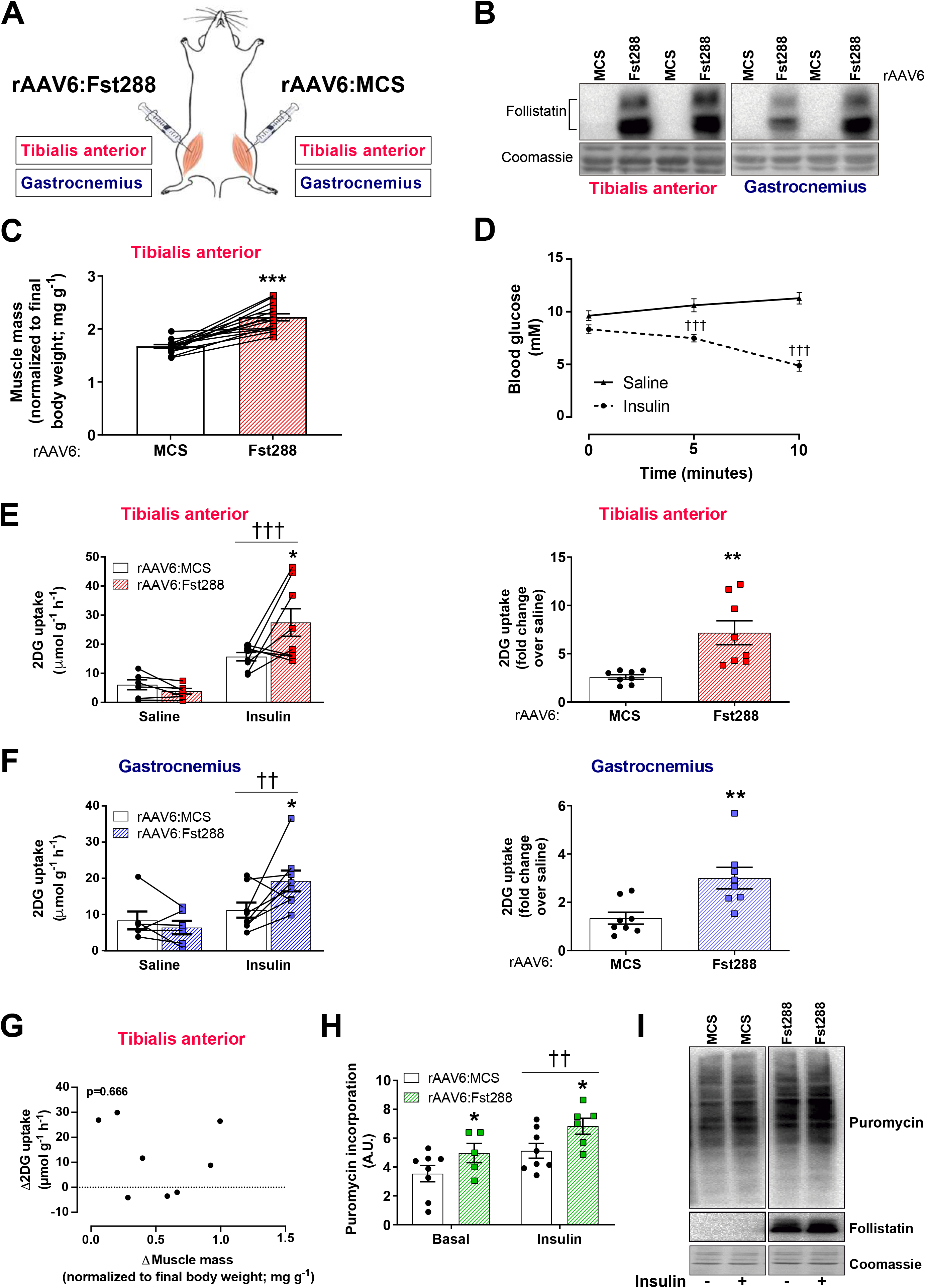
Effect of Follistatin (Fst288) overexpression on muscle glucose uptake. **A)**Illustration of experimental set-up: recombinant adeno-associated virus encoding Fst288 (rAAV6:Fst288) was injected into tibialis anterior and gastrocnemius muscles of one leg, while the contralateral muscles were injected with rAAV6:MSC as control. **B)**Representative blot of follistatin in muscle. **C)**Muscle mass of tibialis anterior (n=14). **D)**Blood glucose levels measured before (0 min) and 5 min and 10 min following retro-orbital insulin (0.3 U kg^−1^ body weight) or saline injection 2 weeks after rAAV6:Fst288 administration in mice (n=6-8). Basal and insulin-stimulated 2-deoxy-glucose (2DG) uptake and fold-change (over saline) in **E)**tibialis anterior and **F)**gastrocnemius muscles from mice 2 weeks after intramuscular rAAV6:Fst288 administration (n=6-8). **G)**Correlation between the change in muscle mass and insulin-stimulated glucose uptake in tibialis anterior muscle weeks after rAAV6:Fst288 administration (n=8). **H)**Insulin-stimulated (10 nM, 60 min) puromycin incorporation in L6-GLUT4myc myotubes following rAAV6:Fst288 administration or as a control rAAV6:MCS. Each experiment was assayed in duplicates or quadruplicates and repeated three times (one sample was excluded due to poor quality of the cells in the well). **I)**Representative blots. Effect of insulin-stimulation is indicated by ††/††† *p* <0.01/0.001. Statistically significant effect of Fst288 overexpression is indicated by */**/\*\*\**p* <0.05/0.01/0.001. Data are shown mean ± SEM with individual values shown when applicable.

### Fst288 overexpression augments insulin-stimulated AKT and PAK1 signaling in muscle

As insulin stimulated, but not basal, glucose uptake was markedly increased independently of the hypertrophy magnitude, we next sought to determine the molecular explanation for this and used immunoblotting techniques to analyze intracellular insulin signaling as illustrated in Fig. 2A. Overexpression of Fst288 significantly upregulated insulin-stimulated p-AKT^Thr308^ (+120%) in TA, but not in gastrocnemius muscles (Fig. 2B). This was also the case for the mTORC2 site, p-AKT^Ser473^ (TA: +50%, Fig. 2C) ^33^. The AKT signaling pathway bifurcates into TBC1D4, important for glucose uptake, and mTORC1, crucial for muscle mass regulation. In line with increased p-AKT in TA, we observed a 68% increase in insulin-stimulated p-TBC1D4^Thr642^ with Fst288 overexpression compared to rAAV6:MCS (Fig. 2D). Like p-AKT, insulin-stimulated TBC1D4^Thr642^ tended to be increased in gastrocnemius with Fst288 overexpression. TBC1D1, a protein with 90% homology to TBC1D4 ^34^, is in muscle phosphorylated at site Thr596 in response to insulin and this is likely downstream of Akt ^35–37^. However, surprisingly, Fst288 overexpression lowered both basal and insulin-stimulated p-TBC1D1^T596^ in TA (basal: −44%; insulin: −41%) and gastrocnemius muscle (basal: −43%; insulin: −39%) (Fig. 2E). Fst288 overexpression resulted in a remarkable increase (+400%) in insulin-stimulated (but not basal) p-p70S6K^Thr389^ (Fig. 2F), an endogenous substrate of mTORC1, suggesting that Fst288 overexpression increases anabolic insulin action in mature skeletal muscle. A less described, but likely equally important signalling pathway for insulin action, is the Rac1-mediated pathway. Rac1’s downstream target, PAK1 has been identified by proteomic muscle profiling to be a putative target for Fst ^19,38^, although actual activation through this pathway has not been investigated. Indeed, insulin-stimulated phosphorylation of PAK1 at its activation site Thr423, was upregulated by rAAV6:Fst288 (TA: +56%, gastrocnemius: +29%) compared to rAAV6:MCS control (Fig. 2G). Thus, Fst288 augments signaling in skeletal muscle to increase both glucose uptake and muscle mass in response to insulin. Except for p-TBC1D1^Thr596^, no effect of Fst288 was observed on any of the analyzed signaling proteins during saline treatment, suggesting that the molecular modifications in this signaling pathway in response to Fst depend on the stimulation with insulin.

**Figure 2.**
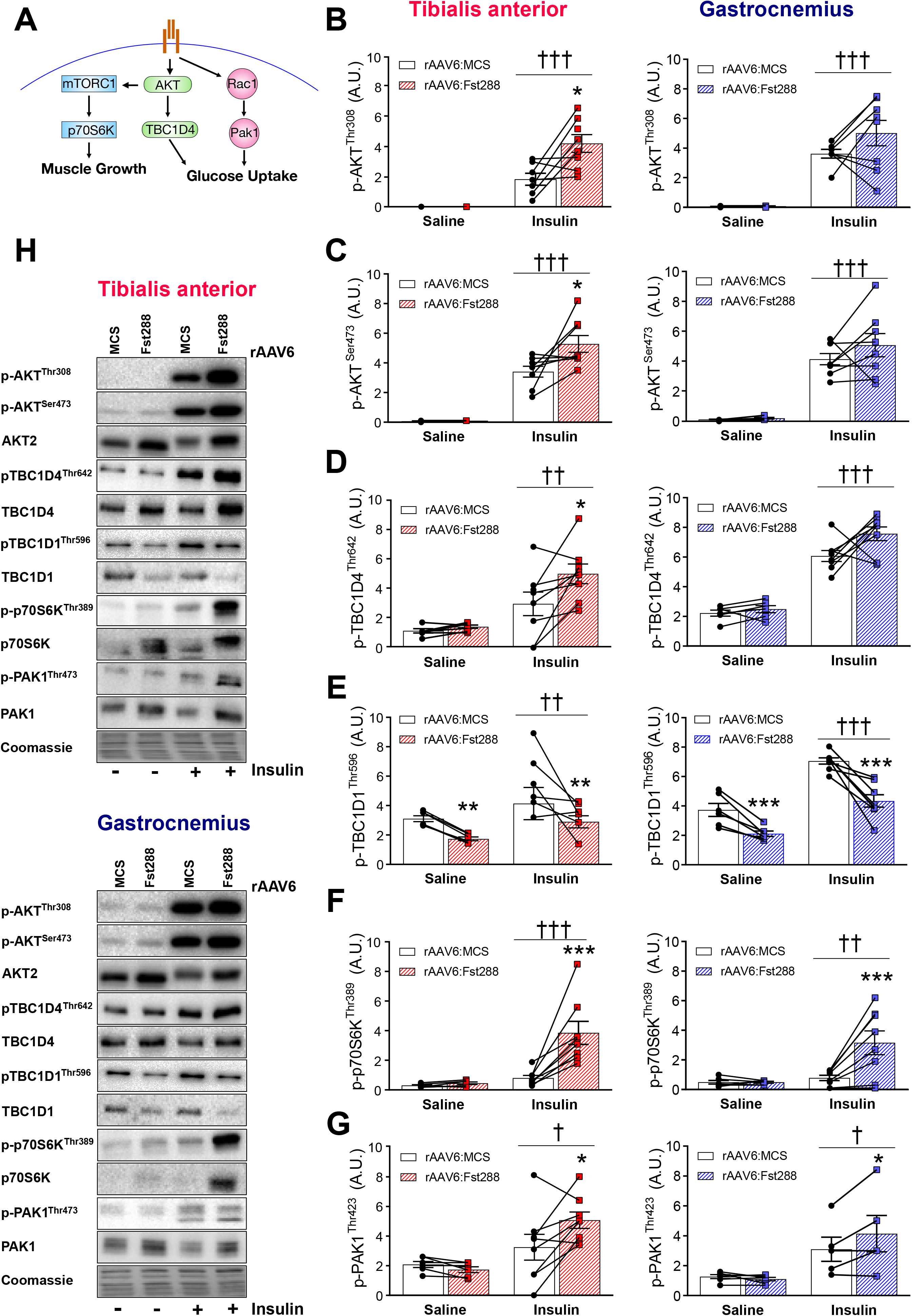
Insulin-stimulated intracellular signaling in skeletal muscle following rAAV6:Fst288 administration. **A)**Graphic illustration of the signaling pathways analyzed. Basal and insulin-stimulated phosphorylated (p), **B)**p-AKT^Thr308^, **C)**p-AKT^Ser473^, **D)**p-TBC1D4^Thr642^, **E)**p-TBC1D1^Thr596^, **F)**p-p70S6K^Thr389^, and **G)**p-PAK1^Thr423^ in tibialis anterior and gastrocnemius muscles following 2-week rAAV6:Fst288 overexpression in muscle (n=5-8, one to two mouse sample excluded due to poor quality blot). **G)**Representative blots. Effect of insulin-stimulation in Fst288 groups or main effect of insulin stimulation is indicated by ††/††† 0070 <0.01/0.001. Statistically significant effect of Fst288 overexpression on insulin signaling is indicated by */**/\*\*\**p* <0/0.05/0.01/0.001. Data are shown as mean ± SEM with individual values.

### Fst288 overexpression increases muscle content of key insulin sensitive signaling proteins but does not affect content of proteins involved in glucose handling

We next sought to investigate whether the increase in insulin sensitivity could be due to an increased capacity of the muscles to enhance insulin-mediated molecular signals. We therefore analyzed the total content of proteins involved in insulin signal transduction and glucose handling. As expected, we observed no difference on protein expression between the 10 minutes saline or insulin-stimulated conditions (see representative blots Fig. 3), and thus they were pooled to increase statistical power and accuracy. In TA muscle, we found a marked upregulation of AKT2 by rAAV6:Fst288 (+90%, Fig. 3A) and TBC1D4 (+60%, Fig. 3B). In contrast, TBC1D1was downregulated to half of the rAAV6:MCS treated control muscles (Fig. 3C). Similar to AKT2 and TBC1D4, total p70S6K protein tripled (Fig. 3D), and PAK1 protein was doubled (Fig. 3E). Because the levels of AKT2, TBC1D4, and p70S6K increased, the ratio of p-protein/total protein was similar for rAAV6:Fst288 and rAAV6:MCS administered muscles (Supplemental Fig. 1), suggesting that Fst288 overexpression increased protein expression and activity in the insulin signaling pathway to a similar extent. In contrast, the ratio of p-protein/total protein was decreased for PAK1 with Fst288 (Supplemental Fig. 1). Due to the reduced protein expression, we only observed minor effects when relating p-TBC1D1^Thr596^ to the total protein content (Supplemental Fig. 1). In addition to the above mentioned proteins involved in insulin signal transduction, insulin receptor substrate 1 (IRS1) was also upregulated (+33%) in response to Fst288 overexpression (Fig. 3F). In contrast, protein content of the glucose handling machinery e.g. GLUT4 (glucose transporter) and hexokinase (a proposed rate-limiting enzyme to *in vivo* glucose uptake in skeletal muscle ^39^), was not regulated by Fst288 (Fig. 3G+H). We also analyzed complexes of the electron transport chain and found that all the investigated complexes were unaffected by Fst288 (Fig. 3I). We found no effect in the phosphorylated levels of the cell stress sensor AMP-activated protein kinase (AMPK) (Supplemental Fig. 1), suggesting that Fst did not alter intracellular metabolic stress. We observed similar regulation of proteins in gastrocnemius muscle (data not shown), except for the ratio of p-protein/total protein for p-AKT^Ser473^ which was significantly decreased with Fst288 (Supplemental Fig. 1). These findings show that Fst288 alters the expression of key insulin sensitive proteins but does not regulate proteins directly involved in glucose handling.

**Figure 3.**
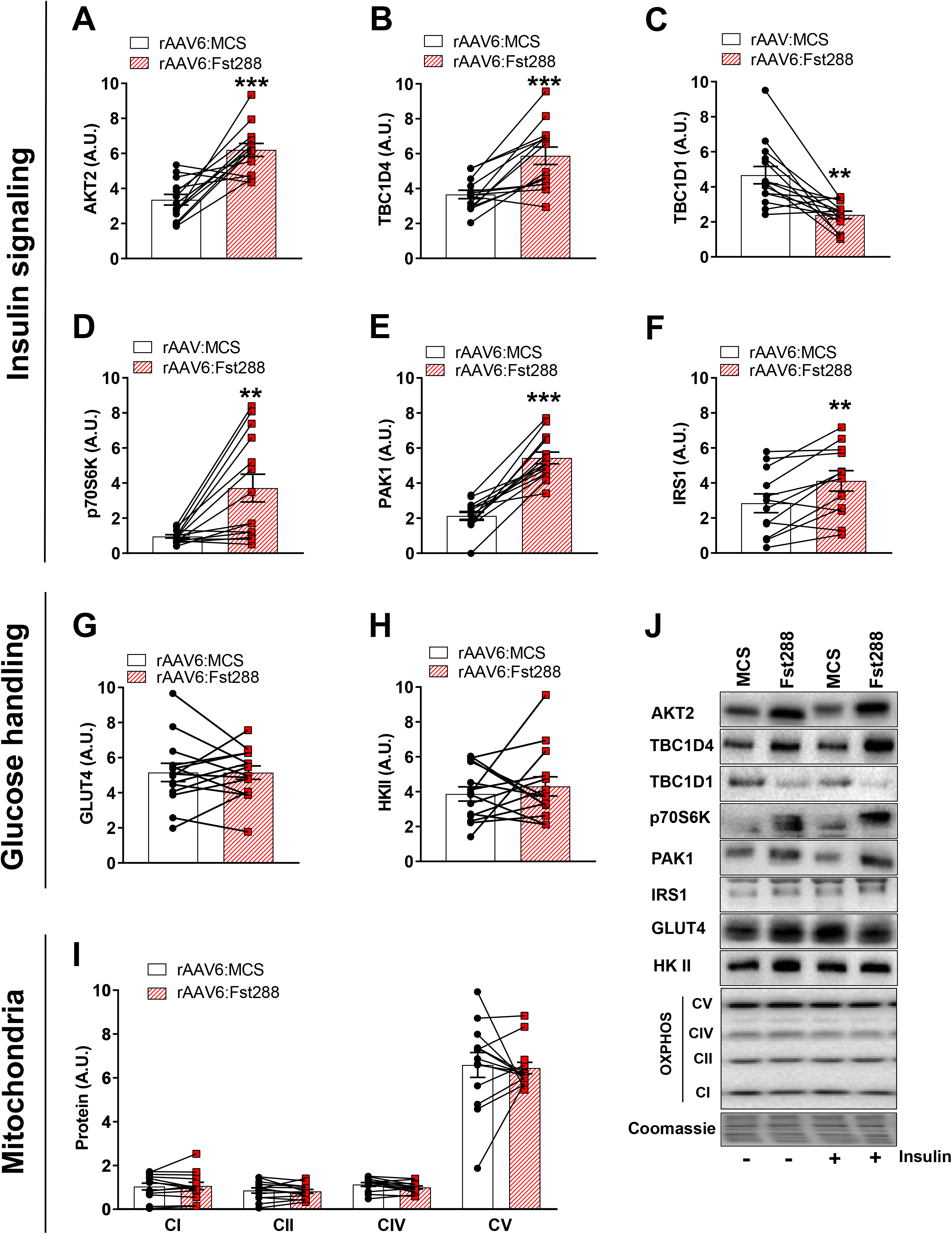
Effect of Fst288 overexpression on proteins involved in insulin signaling and glucose handling. Protein expression of **A)**AKT2, **B)**TBC1D4, **C)**TBC1D1, **D)**p70S6K, **E)**PAK1, **F)**IRS1 **G)**GLUT4, **H)**HKII, and **I)**complexes of the electron transport chain in tibialis anterior muscle following 2-week rAAV6:Fst288 overexpression in muscle. No difference was observed in protein expression between the 10 minutes saline and insulin-stimulated conditions, and thus they were pooled to increase statistical power and accuracy of effect-size estimation (n=12-14, one to two mouse samples excluded due to poor quality blot). **J)**Representative blots. Statistically significant effect of Fst288 overexpression is indicated by **/\*\*\**p* <0.01/0.001. Data are shown as mean ± SEM with individual values.

### Fst288 overexpression increases muscle content and activity of PDH-E1α in gastrocnemius muscle

Having found that Fst288 overexpression increased glucose uptake in muscle, we next asked what the metabolic fate of the extra glucose taken up was. The pyruvate dehydrogenase (PDH) complex catalyzes the irreversible conversion of pyruvate to acetyl CoA and phosphorylation of site 1, p-PDH-E1α^S293^ (basal: −13%; insulin: −20%, Fig. 4A) and site 2, p-PDK-E1α^S300^ (basal: −23%; insulin: −42%, Fig. 4B) were downregulated by Fst288 in gastrocnemius but not TA muscle, suggesting increased activity of the PDH complex ^40^. The level of the PDH-E1α subunit was unaffected by Fst288 in TA muscle, but decreased (−13%) in gastrocnemius (Fig. 4C). Therefore, p-protein/total protein was similar for rAAV6:Fst288 and rAAV6:MCS administered muscles for site 1 and only slightly downregulated for site 2 (basal: −11%; insulin: −36%, Supplemental Fig. 2). Levels of PDH kinase 4 (PDK4), a protein that phosphorylates and inactivates PDH ^40–42^, were unchanged in both TA and gastrocnemius muscle in response to Fst288 overexpression (Fig. 4D). Together, these findings suggest that Fst288 increased the capacity for glycolysis-driven mitochondrial glucose oxidation in gastrocnemius muscle, thereby increasing the reliance on carbohydrate metabolism. *Fst288 overexpression restores insulin resistance in diet-induced obese mice.* Since we observed that Fst markedly increased insulin-stimulated glucose uptake in skeletal muscle, we next asked if Fst288 could be relevant for restoring insulin resistant conditions. We fed mice a 60E% high-fat diet (HFD) for 10 weeks, an intervention known to induce obesity and muscular insulin resistance ^43^. Mice were injected with rAAV6: Fst288 eight weeks into the diet intervention, at a time point when the HFD mice weighed significantly more than chow fed control mice (Fig. 5A), displayed markedly increased adiposity (Fig. 5B), and reduced relative lean body mass (Fig. 5C). Importantly, at this time of the HFD intervention, muscular insulin resistance has already set in ^44,45^, thus allowing us to test the therapeutic potential for Fst288 in restoring insulin sensitivity in already insulin resistant muscle. That the mice were indeed insulin resistant on the experimental day was evidenced by a blunted blood glucose response to insulin (dose 0.3U kg^−1^) in the HFD fed mice compared with chow (Fig. 5D) ^46,47^. We verified that TA muscle mass was increased by 33% after 2 weeks (Fig. 5E), and that Fst was expressed in the rAAV6:Fst288 treated muscles (Fig. 5F). The HFD intervention did not alter basal or insulin-induced intracellular signaling judged by p-AKT^T308^ and p-AKT^S473^ (data not shown) in agreement with some other reports ^26,48^. Basal muscle glucose uptake in HFD mice was unaffected by diet and Fst288 administration (Fig. 5G). HFD induced a 40% reduction in insulin-stimulated glucose uptake in TA and gastrocnemius muscles (Fig. 5H) when comparing with the rAAV6:MCS muscles from chow fed mice. Remarkably, Fst288 treatment completely restored insulin-stimulated glucose uptake (Fig. 5H,I,J) in the insulin resistant muscles of HFD fed mice. Having found that Fst288 overexpression increased glucose uptake in muscle and could restore insulin action, we next measured muscle glycogen. In both insulin sensitive chow fed and insulin resistant HFD fed mice, Fst288-treated TA muscles contained significantly more glycogen (+45%), compared to controls (Fig. 5K). These findings suggest that the larger amount of glucose that is taken up by the rAAV6:Fst288-administered muscles is, at least in part, incorporated into glycogen. Together, our findings suggest that Fst treatment could be a potent strategy to increase insulin sensitivity, also in insulin resistant muscles. A summary of our findings is graphically illustrated in Fig. 6.

**Figure 4.**
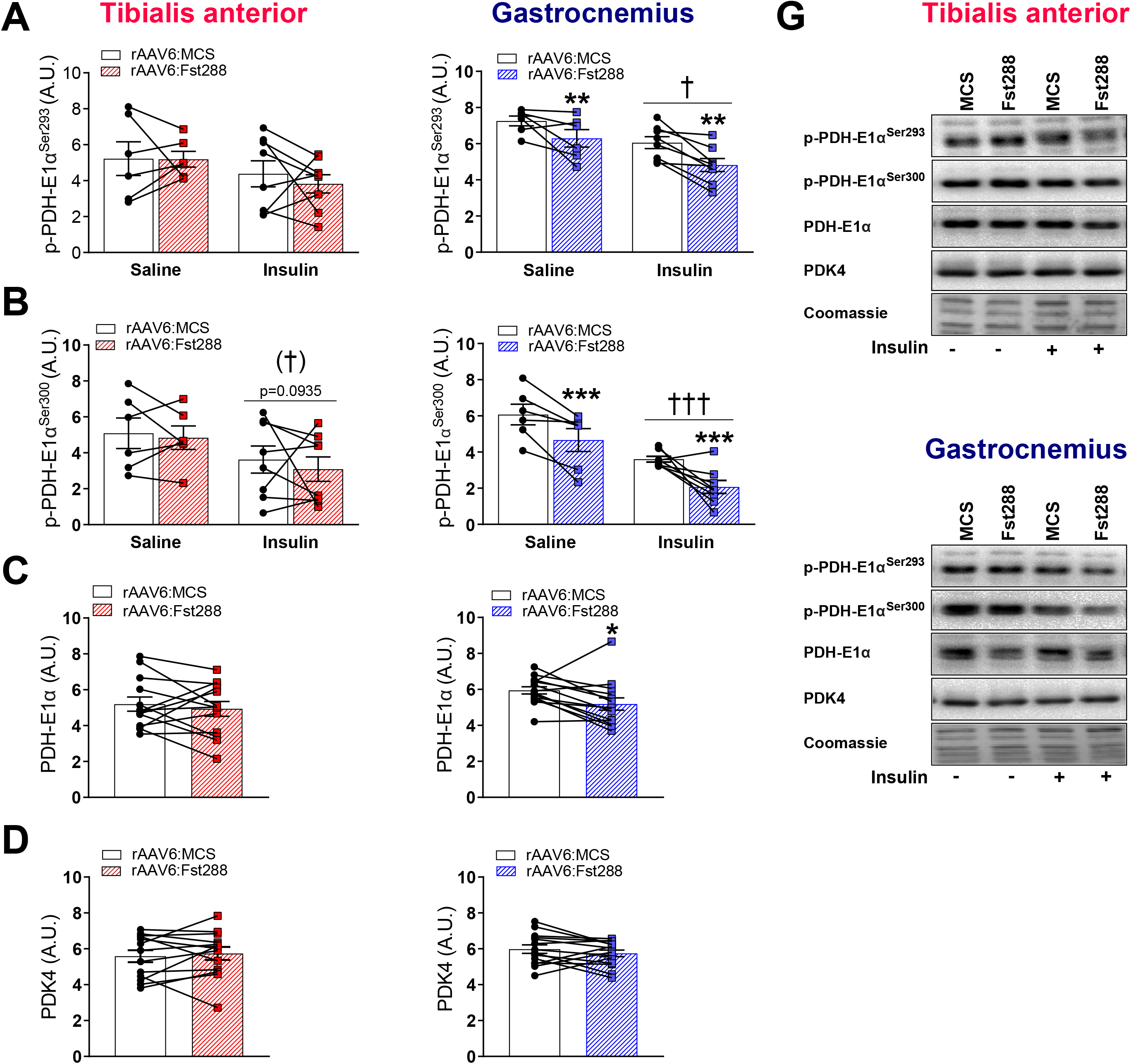
Effect of Fst288 overexpression on PDH activity. Basal and insulin-stimulated phosphorylated (p), **A)**p-PDH-E1α^Ser293^ and **B)**p-PDH-E1α^Ser300^ in tibialis anterior and gastrocnemius muscles following 2-week rAAV6:Fst288 overexpression in muscle (n=6-8). Protein expression of **C)**PDH-E1α and **D)**PDK4 in tibialis anterior and gastrocnemius muscle. No difference was observed in protein expression between the 10 minutes saline or insulin-stimulated conditions, and thus they were pooled to increase statistical power and accuracy (n=13-14, out of lysate for one tibialis anterior muscle sample). **E)**Representative blots of puromycin and follistatin in L6-GLUTmyc myotubes with or without insulin-stimulation following rAAV6:administration. Main effect of insulin-stimulation is indicated by †/††† *p* <0.05/0.001. Statistically significant main effect of Fst288 overexpression is indicated by */**/\*\*\**p* <0.05/0.01/0.001. Data are shown as mean ± SEM with individual values.

**Figure 5.**
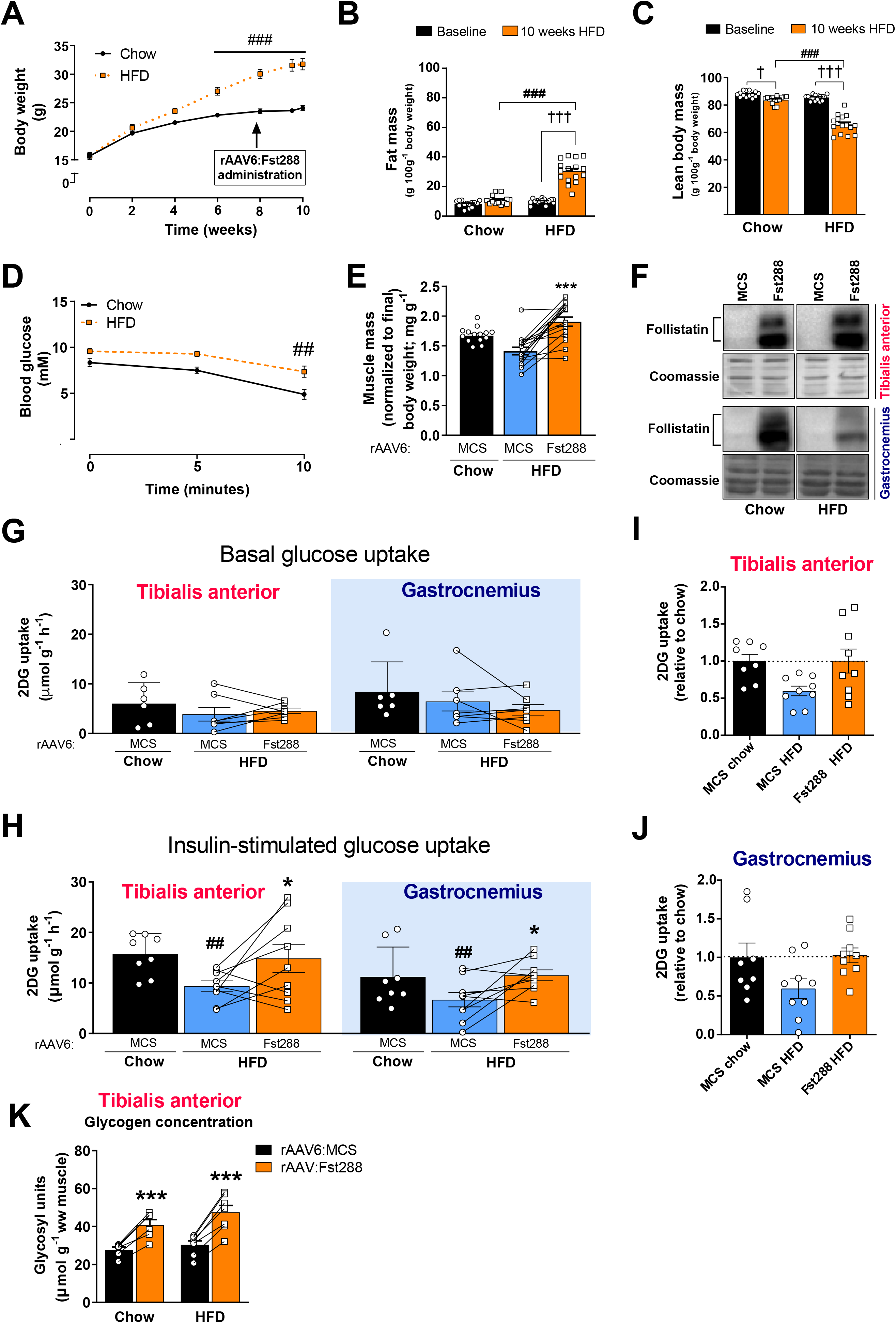
Effect of Fst288 overexpression in insulin-resistant obese mice. **A)**Body weight, **B)**fat mass, and **C)**lean body mass of mice fed a standard chow diet or 60E% high-fat diet (HFD) for 10 weeks (n=14–16). **D)**Blood glucose in response to insulin (retro-orbitally injected, 0.3 U kg^−1^ body weight) (n=8-9). **E)**Muscle mass of tibialis anterior muscle (n=14-16). **F)**Representative blots of follistatin in tibialis anterior and gastrocnemius muscle 2 weeks following intramuscular rAAV6:Fst288 injection. **G)**Basal (saline-injected) and **H)**insulin-stimulated 2 deoxy-glucose (2DG) uptake in tibialis and gastrocnemius muscles intramuscularly injected with rAAV6:Fst288 or rAAV6:MCS (control). Fold change of insulin-stimulated 2DG uptake in HFD over chow rAAV6:MCS in **I)**tibialis anterior and **J)**gastrocnemius muscle. **K)**Glycogen content determined as glycosyl units in tibialis anterior muscle (n=6-8). Statistical significant difference between baseline and 10-week time point is indicated by †/††† *p* <0.05/0.001.Statistically significant effect of Fst288 overexpression is indicated by */\*\*\**p* <0.05/0.001. Statistical significance between HFD and chow control mice is indicated by ##/### *p* <0.01/0.001. Data are shown mean ± SEM with individual values shown when applicable.

**Figure 6.**
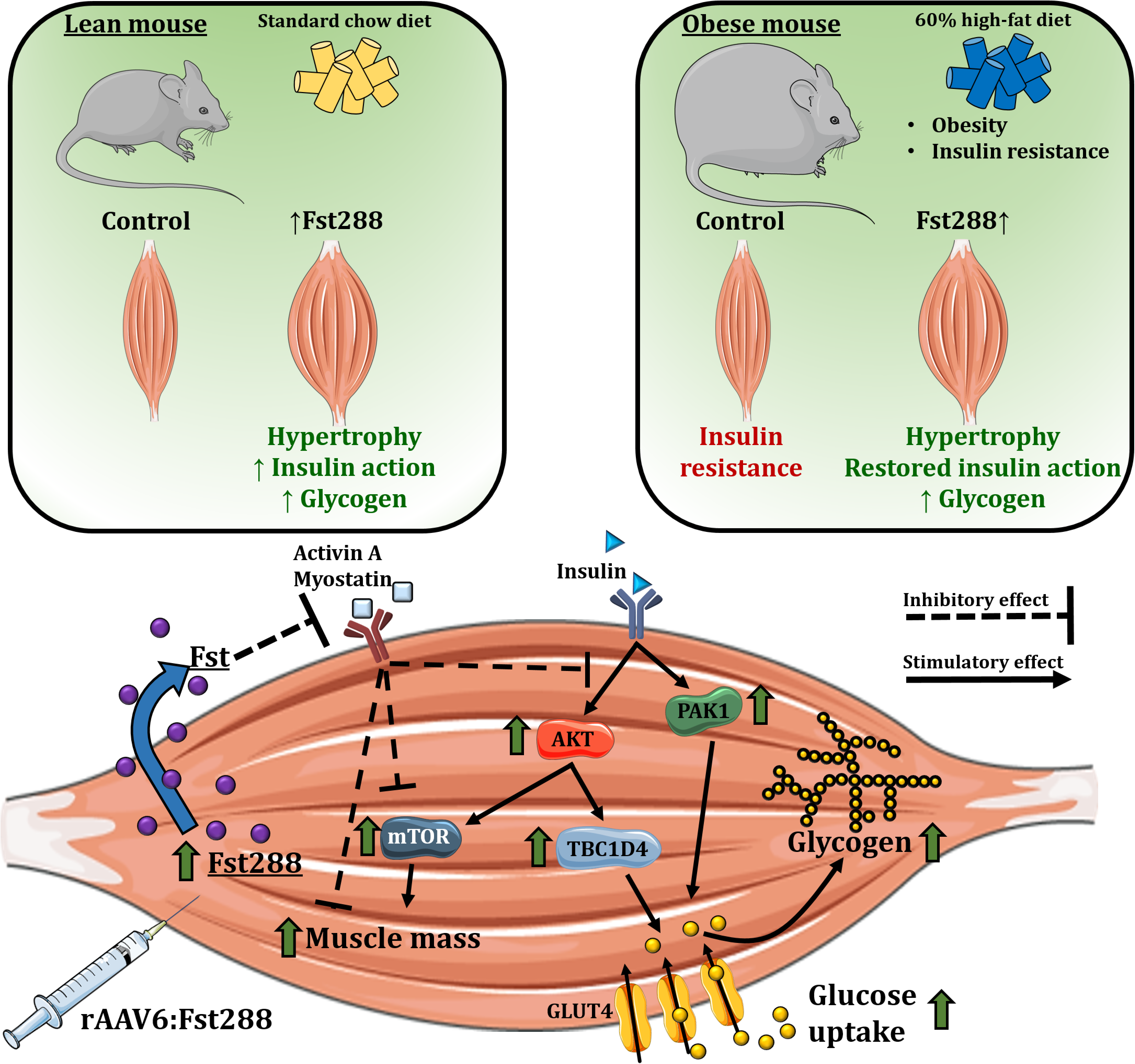
A graphical illustration of our main findings. Fst288 muscle overexpression markedly increased insulin-stimulated (but not basal) intracellular insulin signalling of AKT, TBC1D4, PAK1, and mTOR signalling via p70S6K. In lean insulin sensitive mice, muscle-directed Fst288 overexpression increased skeletal muscle insulin-stimulated glucose uptake. Fst288 completely normalized muscle glucose uptake in insulin-resistant obese mice.

### Bariatric surgery increases Fst and reduces activin A in obese insulin resistant individuals

Having established that Fst regulates insulin action in skeletal muscle, we next investigated the relationship between insulin action and TGF-β ligands in humans. We measured circulating Fst and activin A concentrations in a group of obese insulin resistant (but normoglycemic) individuals before, 1 week, and 1 year after treatment by Roux-en-Y gastric bypass surgery (RYGB). This procedure has consistently been shown to produce marked and rapid improvements in insulin resistance and glycemic control and data for the currently studied subjects has been published previously^29^. One week following RYGB circulating Fst drastically increased (+108%, Fig. 7A) while circulating activin A was mildly reduced (−26%, Fig. 7B). At the same time hepatic insulin sensitivity and plasma glucagon nearly doubled, while the peripheral insulin resistance did not differ from before surgery ^29,49^. Importantly, the ratio between Fst and activin A (which determines actual activin A signaling through the ActRII), was 217% increased 1 week post RYGB (Fig. 7C) where weight loss has not yet set in (Fig. 7D). These levels were largely maintained 1 year post surgery (Fst +111%; activin A +13%; ratio +168%) when peripheral insulin sensitivity was improved (+200%), and hepatic insulin sensitivity was further improved (+50 %) compared with 1 week after surgery ^29^. These findings indicate that in humans circulating Fst is markedly altered by metabolic status beyond the effect of body weight loss and could thus play an unrecognized critical role in maintenance of insulin sensitivity, glucose homeostasis and muscle mass.

**Figure 7.**
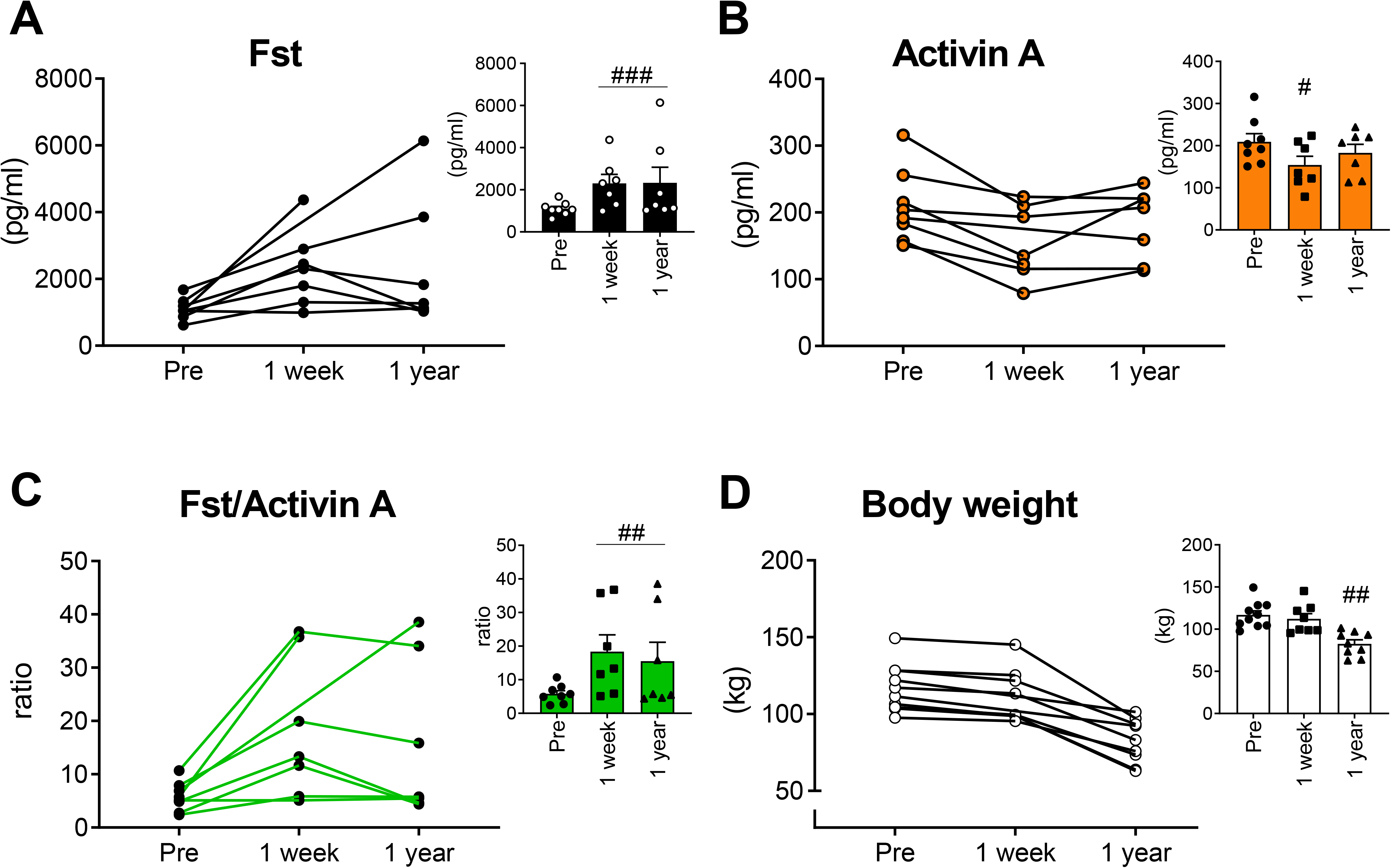
Serum follistatin and activin A concentrations in obese insulin resistant individuals before and following Roux-en-Y gastric bypass surgery. Serum concentrations of **A)**follistatin (Fst), **B)**activin A, **C)**ratio between Fst and activin A before and 1 week and 1 year after Roux-en-Y gastric bypass surgery (RYGB), a procedure that markedly improves glycemic control. **D)**Body weight (n=9, however for 2 data points a value is missing, as indicated in the graph showing individual values). Statistically significant effect of RYGB is indicated by #/##/###*p* <0.05/0.001/0.0001. Data are shown mean ± SEM and/or individual values when applicable.

## Discussion

A major discovery in the current study was that concomitantly with increased muscle mass, Fst288 overexpression markedly augmented insulin action on glucose uptake and intracellular signaling and completely restored insulin resistance in diet-induced obese mouse muscles. To the best of our knowledge, we are the first to describe and mechanistically explain this insulin sensitizing effect of Fst in muscle. This is an important discovery, as muscle wasting conditions are often associated with muscular insulin resistance, including in cancer cachexia, sarcopenia, and Duchenne muscular dystrophy. Thus, strategies to treat insulin resistance in muscle wasting diseases are warranted and our current study suggests that Fst might be an attractive candidate. Our findings are supported by a recent study using a transgenic (TG) approach to overexpress Fst in the entire skeletal muscle mass ^50^. In that study, Fst-TG mice displayed better glucose tolerance compared with control mice, however tissue-specific insulin action was not investigated. Other circumstantial evidence supports our results that Fst is a positive regulator of insulin sensitivity in muscle. For example, pharmacological inhibition or genetic deletion of the Fst targets, myostatin or activin A, protects against HFD-induced obesity ^51–53^. Such findings support our results that Fst is a positive regulator of insulin sensitivity in muscle. Importantly, the role for this pathway in insulin action is also evident from clinical trials in humans. Here, the ActRII inhibitor bimagrumab markedly improved insulin sensitivity and HbA1c in insulin resistant individuals ^11^. None of the above studies were able to ascribe the increased insulin sensitivity to a specific tissue, but our results suggest that likely, improved skeletal muscle glucose uptake underlies, at least some of, the beneficial effects of inhibiting ActRII or its ligands. Additionally, our data indicate that Fst *per se* and not the magnitude of hypertrophy drives the increase in insulin action placing Fst as a central mediator in the crosstalk between insulin action and muscle wasting. Intriguingly, the above mentioned evidence and our results appear to contradict a recent study showing that when Fst was overexpressed in the liver, insulin sensitivity and glucose tolerance was impaired in mice with genetically-induced liver insulin resistance ^55^. However, in that study viral liver-expression of Fst markedly altered RNA expression profiles of the liver, probably affecting insulin sensitivity by confounding unknown factors. Furthermore, Fst was not overexpressed in skeletal muscle. Our data firmly show that Fst288 overexpression in muscle improves insulin action in a set-up where the effect can be isolated to muscular effects, as the Fst288 variant used in our muscle study typically remains localized in the immediate vicinity of the cell from which it is secreted ^56^. This is in contrast to the longer circulating Fst315 isoform. Clearly, future work should investigate the tissue-specific metabolic effects of distinct Fst isoforms.

Another major finding of our study was the marked upregulation of circulating Fst in obese insulin resistant humans following RYGB surgery, a treatment that markedly improved whole body insulin sensitivity ^29,49^. Furthermore, the TGF-β ligand, activin A was reduced following RYGB. As a result, the Fst/activin A ratio increased, suggesting a marked reduction of the ActRII ligand, activin A available to activate TGB-β signaling. However, peripheral insulin sensitivity was not yet improved 1 week after RYGB where an increase in Fst and decline in activin A was already observed. Our results may have broader implications also to adipose tissue function, as Fst can induce browning ^57^, which could promote increased energy expenditure and thereby protect from obesity. While our findings show that circulating Fst is markedly upregulated in humans by a treatment that improves glucose homeostasis, future studies should confirm inactivation of ActRII signaling in muscle biopsy samples. The mechanisms behind the upregulation of Fst in humans following RYGB are unresolved, but prolonged fasting (72 h) in humans has been demonstrated to increase circulating Fst 155% and decrease activin A ^58^, mimicking the results from the current study. Additionally, glucagon is suggested to increase the secretion of Fst from the liver ^59^. As such, the effect 1 week following RYGB could be a result of the dramatic reduction in food intake by these patients and a concomitant increase in plasma glucagon in the first week following surgery, while the mechanism for the upregulation 1 year following RYGB surgery remains unknown. In humans, the liver is a key contributor to circulating Fst, which is regulated by the glucagon-to-insulin ratio ^59^, further suggesting that circulating Fst is closely related to energy metabolism. However, contrasting with Fst’s proposed beneficial role in insulin sensitivity, plasma Fst is actually slightly elevated in patients with type 2 diabetes ^60^ and in nonalcoholic fatty liver disease ^61^. Those studies did not analyze activin A. Thus the Fst/activin A ratio could not be determined, which would be the relevant read-out for actual activation/inactivation of ActRII. Thus, the exact cause(s) and consequence(s) of circulating Fst remain to be determined. It is worth noting that activin A is not the only ligand of Fst and thus the effect of Fst could be due to inhibition of other ligands like myostatin. Indeed, insulin-stimulated glucose uptake is slightly increased in skeletal muscle of myostatin knockout (KO) mice ^52^. However, Fst overexpression induced muscle hypertrophy to a similar extent in myostatin KO and control mouse muscle ^62^, suggesting that myostatin is dispensable for Fst to induce hypertrophy. Future studies should determine the role of myostatin in Fst-induced enhanced insulin sensitivity.

The molecular mechanism by which Fst regulates muscular insulin sensitivity was investigated in our study. Fst288 overexpression caused upregulation of key mediators of glucose uptake, including, AKT, TBC1D4, and PAK1 and enhanced their responsiveness to insulin. Furthermore, the insulin-responsiveness of p-p70S6K, a reporter of mTORC1 activity, was increased by 4-fold. Seminal studies have shown that phosphorylated AKT induces mTOR-mediated phosphorylation of p70S6K in skeletal muscle ^63^, resulting in muscle hypertrophy ^64^. The increased insulin-stimulated p-p70S6K in gastrocnemius with Fst288 overexpression even though p-AKT was not significantly increased could be explained by signal amplification downstream of Akt, as p70S6K activity previously has been shown to be preserved in mice muscle with severe insulin receptor defects ^65^, suggesting either signal amplification or another upstream kinase regulating p70S6K in response to insulin. Although p70S6K seems to be largely dispensable for Fst-induced hypertrophy ^12,66^, our study shows that Fst288 might also augments insulins action on anabolic signals that regulate muscle mass. Indeed, both basal and insulin-stimulated puromycin incorporation in L6 myotubes was upregulated by Fst overexpression, suggesting increased protein synthesis. This could be relevant as a blunted protein anabolic response to insulin has been proposed in obesity ^67^ and T2D ^7^, although this was not directly tested in our study. Interestingly, despite an upregulated total amount of proteins in the insulin signaling pathways, we observed no effect of Fst288 on basal intracellular signaling or basal glucose uptake. This suggests that, while Fst increases the capacity for insulin signaling to be activated, insulin is needed to make use of this capacity. Being able to turn this pathway off in the presence of Fst is likely essential to maintain cellular homeostasis, as constitutive activation of Akt-mTOR would result in activation of negative feedback loops with proven adverse effects on muscle ^68^. Whether Fst could improve skeletal muscle protein synthesis in situations of anabolic insulin resistance often found in ageing ^6^, would be important to determine and could have implications for the progression of sarcopenia in older subjects.

*In vivo* glucose uptake by skeletal muscle is regulated by delivery, transport across the muscle cell surface by the glucose transport GLUT4 and intramyocellular metabolism^69^. In the present study, GLUT4 expression was unchanged with Fst288, while the intracellular signals to translocate GLUT4 to the plasma membrane were greatly enhanced. In addition, HK levels were unaffected, while we observed that Fst increased PDH activity in gastrocnemius muscle. So, in addition to the effects of Fst on key insulin sensitive proteins and glucose incorporation into glycogen, Fst also increased PDH activity in gastrocnemius muscle, suggesting greater capacity for glycolysis-driven mitochondrial glucose oxidation ^52^. This is in line with a previous report showing increased respiratory exchange rate in myostatin KO mice ^52^.Interestingly, we observed that Fst288 regulated the Rab GAP proteins in opposite directions, TBC1D4 up and TBC1D1 down. These proteins regulate GLUT4 translocation and protein expression ^70^. Downregulation of TBC1D1 would be expected to reduce insulin actin on glucose uptake, as TCB1D1 knock-out mouse muscles are insulin resistant, likely due to a 50% reduction in GLUT4 protein ^71^. However on the contrary, reduced TBC1D1 protein and p-TBC1D1^Thr596^ with Fst288 in our study occurred with improved insulin sensitivity and normal GLUT4 protein expression. However, insulin-stimulation promotes phosphorylation of TBC1D1^Thr596^ without affecting the capacity of TBC1D1 to bind to 14-3-3 in L6 cells ^35^ and intact skeletal muscle ^72^, indicating that the decrease in p-TBC1D1^Thr596^ is of minor significance in relation to the upregulated insulin-stimulated glucose uptake with Fst288 overexpression. Interestingly, similar to the increased TBC1D4 phosphorylation observed in our study, some (but not all ^73,74^) studies find that greater post-exercise insulin-stimulated glucose uptake was accompanied by greater TBC1D4 (but not TBC1D1) phosphorylation ^75,76^. Thus, an outstanding question is whether Fst could increase post-exercise insulin sensitivity. Given that our findings show TBC1D4 phosphorylation as a target for Fst, it would be relevant for future studies to elucidate a potential role for Fst in post-exercise enhanced glucose uptake, which also impinges on TBC1D4 ^75,76^.

That Fst markedly affects insulin action in muscle could have significant implications, as Fst treatment and ActRII inhibition are currently under investigation to treat muscle wasting conditions, including Becker muscular dystrophy ^18^ and sarcopenia ^10^. While our study only investigated the effect of short-term Fst288 treatment, clinical trials have found no adverse effects of Fst or pharmacological ActRII inhibition months after gene transfer or treatment in humans ^11,77^. Results from our study suggest that this could be ascribed to the fact that Fst does not affect basal intracellular signaling, while enhancing insulin action. This underlines the therapeutic potential and safety for Fst and ActRII based therapies to improve both muscle mass and as reported in our study, to enhance muscular insulin sensitivity and glucose uptake.

## Acknowledgements

We acknowledge the skilled technical assistance of Irene Bech Nielsen, Betina Bolmgren and Nicoline Resen Andersen (Dept. of Nutrition, Exercise and Sports, Copenhagen University) and we thank all the individuals who took part in these studies. We thank Steffen H. Raun for help with the graphical abstract. We also thank Professor Amira Klip (Cell Biology Program, Research Institute, The Hospital for Sick Children, Toronto, Canada) for the kind donation of the L6-GLUT4myc muscle cells and Professor D. Grahame Hardie (University of Dundee, Scotland, UK) for the kind donation of the PDH-1Eα, p-PDH-1Eα^Ser293^, p-PDH-1Eα^Ser300^ and PDK4 antibodies for this study. The authors certify that they comply with the ethical guidelines for authorship and publishing of the Journal of Cachexia, Sarcopenia and Muscle ^78^.

## Author contributions

X.H., L.L.V.M., E.A.R, and L.S. designed the study. X.H., L.L.V.M., E.D.G., and L.S., designed and conducted the mouse study, with experimental contribution from T.E.J, C.H.O, Z.L., J.R.K., J.D., and P.G. K.N.B-M and S.M. carried out the human experiments. X.H., L.L.V.M., E.A.R., and L.S. drafted the paper. All authors contributed to the final version of the manuscript.

---

**Supplemental Figure 1.**
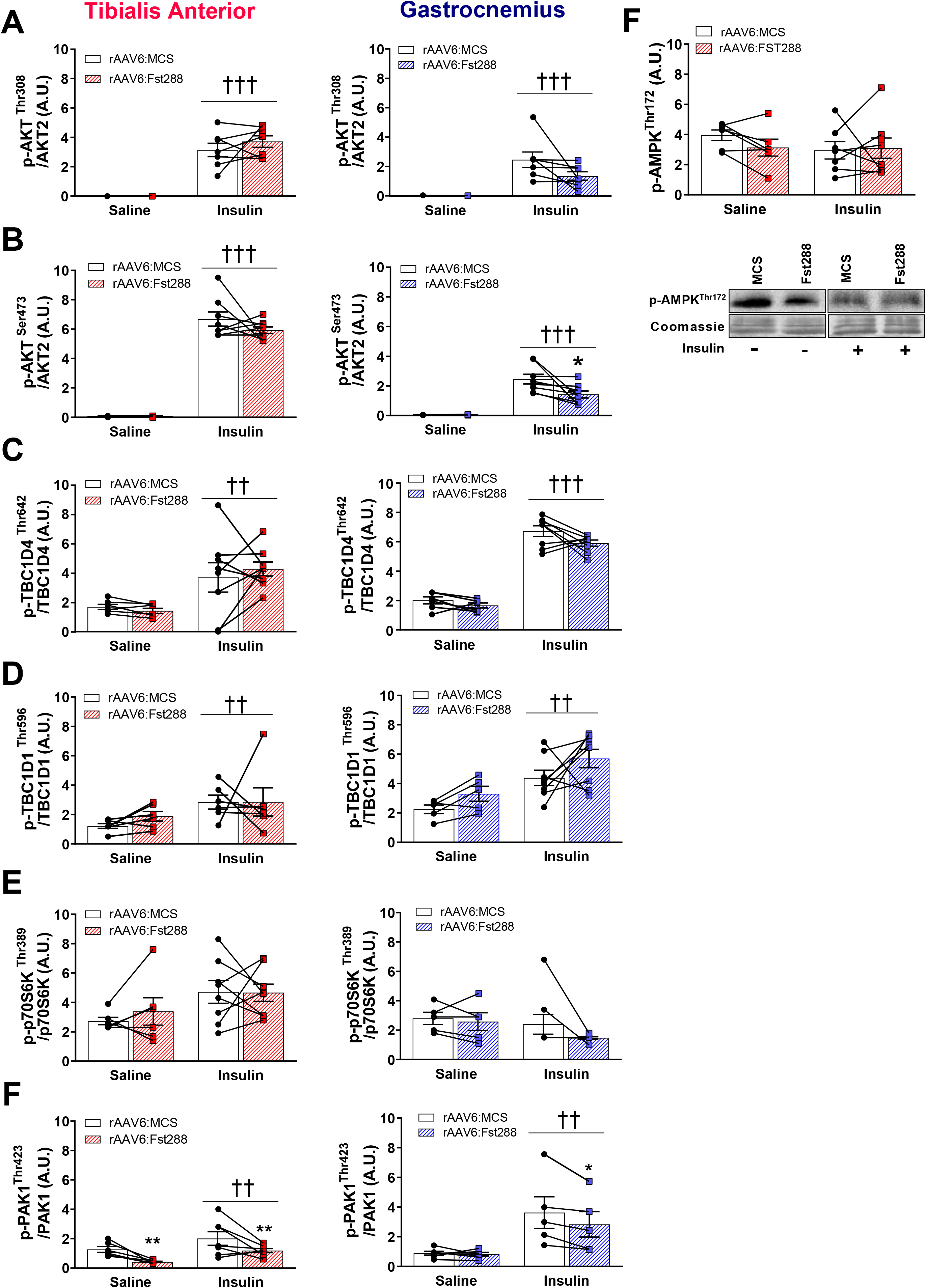
The ratio of phosphorylated (p)-protein/total (t)-protein **A)**of AKT^Thr308^ and **B)** AKT^Ser473^, **C)**TBC1D4^Thr642^, **D)**TBC1D1^Thr596^, **E)**p70S6K^Thr389^, and **F)**PAK1^Thr423^ in tibialis anterior and gastrocnemius muscles following 2-week rAAV6:Fst288 overexpression in muscle (n=5-8, one or two mouse samples excluded due to poor quality blot). **G)**Basal and insulin-stimulated p-AMPK^Thr172^ in tibialis anterior muscle (n=6-8, one mouse sample excluded due to poor quality blot). Main effect of insulin-stimulation is indicated by ††/††† *p* <0.01/0.001. Statistically significant effect of Fst288 overexpression is indicated by */** *p* <0.05/0.01. Data are shown as mean± SEM with individual values.

**Supplemental Figure 2.**
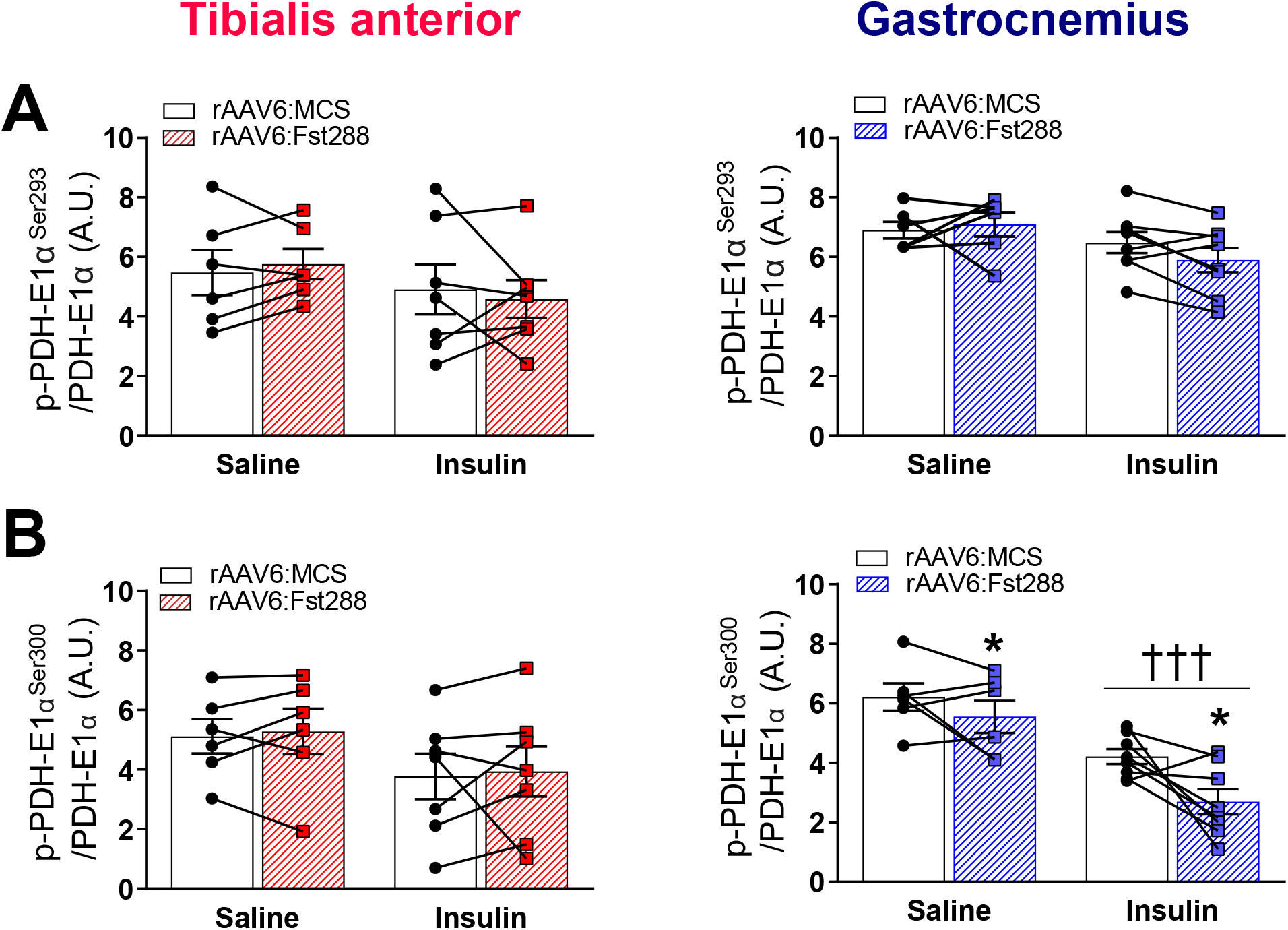
The ratio of phosphorylated protein/total protein of **A)**p-PDH-E1α^Ser293^ and B)p-PDH-E1α^Ser300^ in tibialis anterior and gastrocnemius muscle following 2-week rAAV6:Fst288 overexpression in muscle (n=6-8, out of lysate for one tibialis anterior muscle sample). Main effect of insulin-stimulation is indicated by ††† *p* <0.001. Statistically significant effect of Fst288 overexpression is indicated by \**p* <0.05. Data are shown as mean± SEM with individual values.

## References

1. Leong DP, Teo KK, Rangarajan S, Lopez-Jaramillo P, Avezum A, Orlandini A et al. Prognostic value of grip strength: findings from the Prospective Urban Rural Epidemiology (PURE) study. Lancet 2015;386:266–273.

2. DeFronzo RA, Gunnarsson R, Björkman O, Olsson M, Wahren J. Effects of insulin on peripheral and splanchnic glucose metabolism in noninsulin-dependent (type II) diabetes mellitus. J Clin Invest 1985;76:149–155.

3. Dev R, Bruera E, Dalal S. Insulin resistance and body composition in cancer patients. Ann Oncol 2018;29:ii18–ii26.

4. Wagner EF, Petruzzelli M. Cancer metabolism: A waste of insulin interference. Nature 2015;521:430–1.

5. Cleasby ME, Jamieson PM, Atherton PJ. Insulin resistance and sarcopenia: mechanistic links between common co-morbidities. J Endocrinol 2016;229:R67–R81.

6. Rasmussen BB, Fujita S, Wolfe RR, Mittendorfer B, Roy M, Rowe VL et al. Insulin resistance of muscle protein metabolism in aging. FASEB J 2006;20:768–769.

7. Pereira S, Marliss EB, Morais JA, Chevalier S, Gougeon R. Insulin Resistance of Protein Metabolism in Type 2 Diabetes. Diabetes 2008;57:56–63.

8. Cusi K, Maezono K, Osman A, Pendergrass M, Patti ME, Pratipanawatr T et al. Insulin resistance differentially affects the PI 3-kinase- and MAP kinase-mediated signaling in human muscle. J Clin Invest 2000;105:311–320.

9. Latres E, Mastaitis J, Fury W, Miloscio L, Trejos J, Pangilinan J et al. Activin A more prominently regulates muscle mass in primates than does GDF8. Nat Commun 2017;8:15153.

10. Rooks D, Praestgaard J, Hariry S, Laurent D, Petricoul O, Perry RG et al. Treatment of Sarcopenia with Bimagrumab: Results from a Phase II, Randomized, Controlled, Proof-of-Concept Study. J Am Geriatr Soc 2017;65:1988–1995.

11. Garito T, Roubenoff R, Hompesch M, Morrow L, Gomez K, Rooks D et al. Bimagrumab improves body composition and insulin sensitivity in insulin-resistant individuals. Diabetes, Obes Metab 2018;20:94–102.

12. Winbanks CE, Weeks KL, Thomson RE, Sepulveda P V, Beyer C, Qian H et al. Follistatin-mediated skeletal muscle hypertrophy is regulated by Smad3 and mTOR independently of myostatin. J Cell Biol 2012;197:997–1008.

13. Nakatani M, Takehara Y, Sugino H, Matsumoto M, Hashimoto O, Hasegawa Y et al. Transgenic expression of a myostatin inhibitor derived from follistatin increases skeletal muscle mass and ameliorates dystrophic pathology in mdx mice. FASEB J 2008;22:477–487.

14. Kota J, Handy CR, Haidet AM, Montgomery CL, Eagle A, Rodino-Klapac LR et al. Follistatin gene delivery enhances muscle growth and strength in nonhuman primates. Sci Transl Med 2009;1:6ra15.

15. Amthor H, Nicholas G, McKinnell I, Kemp CF, Sharma M, Kambadur R et al. Follistatin complexes Myostatin and antagonises Myostatin-mediated inhibition of myogenesis. Dev Biol 2004;270:19–30.

16. Yaden BC, Croy JE, Wang Y, Wilson JM, Datta-Mannan A, Shetler P et al. Follistatin: A Novel Therapeutic for the Improvement of Muscle Regeneration. J Pharmacol Exp Ther 2014;349:355–371.

17. Sepulveda P V, Lamon S, Hagg A, Thomson RE, Winbanks CE, Qian H et al. Evaluation of follistatin as a therapeutic in models of skeletal muscle atrophy associated with denervation and tenotomy. Sci Rep 2015;5:17535.

18. Mendell JR, Sahenk Z, Al-Zaidy S, Rodino-Klapac LR, Lowes LP, Alfano LN et al. Follistatin Gene Therapy for Sporadic Inclusion Body Myositis Improves Functional Outcomes. Mol Ther 2017;25:870–879.

19. Davey JR, Watt KI, Parker BL, Chaudhuri R, Ryall JG, Cunningham L et al. Integrated expression analysis of muscle hypertrophy identifies Asb2 as a negative regulator of muscle mass. JCI Insight 2016;1.

20. Sylow L, Kleinert M, Pehmøller C, Prats C, Chiu TT, Klip A et al. Akt and Rac1 signaling are jointly required for insulin-stimulated glucose uptake in skeletal muscle and downregulated in insulin resistance. Cell Signal 2014;26:323–331.

21. JeBailey L, Wanono O, Niu W, Roessler J, Rudich A, Klip A. Ceramide- and oxidant-induced insulin resistance involve loss of insulin-dependent Rac-activation and actin remodeling in muscle cells. Diabetes 2007;56:394–403.

22. Sylow L, Jensen TE, Kleinert M, Hojlund K, Kiens B, Wojtaszewski J et al. Rac1 Signaling Is Required for Insulin-Stimulated Glucose Uptake and Is Dysregulated in Insulin-Resistant Murine and Human Skeletal Muscle. Diabetes 2013;62:1865–1875.

23. Blankinship MJ, Gregorevic P, Allen JM, Harper SQ, Harper H, Halbert CL et al. Efficient transduction of skeletal muscle using vectors based on adeno-associated virus serotype 6. Mol Ther 2004;10:671–678.

24. Ferre P, Leturque A, Burnol A-F, Penicaud L, Girard J. A method to quantify glucose utilization in vivo in skeletal muscle and white adipose tissue of the anaesthetized rat. Biochem J 1985;228:103–110.

25. Fueger PT, Hess HS, Posey KA, Bracy DP, Pencek RR, Charron MJ et al. Control of exercise-stimulated muscle glucose uptake by GLUT4 is dependent on glucose phosphorylation capacity in the conscious mouse. J Biol Chem 2004;279:50956–61.

26. Raun SH, Ali M, Kjøbsted R, Møller LL V, Federspiel MA, Richter EA et al. Rac1 muscle knockout exacerbates the detrimental effect of high-fat diet on insulin-stimulated muscle glucose uptake independently of Akt. J Physiol 2018;596:2283–2299.

27. Passonneau JV, Gatfield PD, Schulz DW, Lowry OH. An enzymic method for measurement of glycogen. Anal Biochem 1967;19:315–326.

28. Schmidt EK, Clavarino G, Ceppi M, Pierre P. SUnSET, a nonradioactive method to monitor protein synthesis. Nat Methods 2009;6:275–277.

29. Bojsen-Møller KN, Dirksen C, Jørgensen NB, Jacobsen SH, Serup AK, Albers PH et al. Early enhancements of hepatic and later of peripheral insulin sensitivity combined with increased postprandial insulin secretion contribute to improved glycemic control after Roux-en-Y gastric bypass. Diabetes 2014;63:1725–37.

30. Pilegaard H, Birk JB, Sacchetti M, Mourtzakis M, Hardie DG, Stewart G et al. PDH-E1 Dephosphorylation and Activation in Human Skeletal Muscle During Exercise: Effect of Intralipid Infusion. Diabetes 2006;55:3020–3027.

31. Kiilerich K, Adser H, Jakobsen AH, Pedersen PA, Hardie DG, Wojtaszewski JFP et al. PGC-1α increases PDH content but does not change acute PDH regulation in mouse skeletal muscle. Am J Physiol Integr Comp Physiol 2010;299:R1350–R1359.

32. Welinder C, Ekblad L. Coomassie Staining as Loading Control in Western Blot Analysis. J Proteome Res 2011;10:1416–1419.

33. Sarbassov DD, Guertin DA, Ali SM, Sabatini DM. Phosphorylation and regulation of Akt/PKB by the rictor-mTOR complex. Science 2005;307:1098–101.

34. Cartee GD. Roles of TBC1D1 and TBC1D4 in insulin- and exercise-stimulated glucose transport of skeletal muscle. Diabetologia 2015;58:19–30.

35. Chen S, Murphy J, Toth R, Campbell DG, Morrice NA, Mackintosh C. Complementary regulation of TBC1D1 and AS160 by growth factors, insulin and AMPK activators. Biochem J 2008;409:449–59.

36. Vichaiwong K, Purohit S, An D, Toyoda T, Jessen N, Hirshman MF et al. Contraction regulates site-specific phosphorylation of TBC1D1 in skeletal muscle. Biochem J 2010;431:311–20.

37. Treebak JT, Pehmøller C, Kristensen JM, Kjøbsted R, Birk JB, Schjerling P et al. Acute exercise and physiological insulin induce distinct phosphorylation signatures on TBC1D1 and TBC1D4 proteins in human skeletal muscle. J Physiol 2014;592:351–375.

38. Barbé C, Bray F, Gueugneau M, Devassine S, Lause P, Tokarski C et al. Comparative Proteomic and Transcriptomic Analysis of Follistatin-Induced Skeletal Muscle Hypertrophy. J Proteome Res 2017;16:3477–3490.

39. Fueger PT, Shearer J, Bracy DP, Posey KA, Pencek RR, McGuinness OP et al. Control of muscle glucose uptake: test of the rate-limiting step paradigm in conscious, unrestrained mice. J Physiol 2005;562:925–935.

40. Sugden MC, Holness MJ. Recent advances in mechanisms regulating glucose oxidation at the level of the pyruvate dehydrogenase complex by PDKs. Am J Physiol Metab 2003;284:E855–E862.

41. Roche TE, Baker JC, Yan X, Hiromasa Y, Gong X, Peng T et al. Distinct regulatory properties of pyruvate dehydrogenase kinase and phosphatase isoforms. 2001. pp. 33–75.

42. Patel MS, Korotchkina LG. Regulation of mammalian pyruvate dehydrogenase complex by phosphorylation: complexity of multiple phosphorylation sites and kinases. Exp Mol Med 2001;33:191–197.

43. Kleinert M, Clemmensen C, Hofmann SM, Moore MC, Renner S, Woods SC et al. Animal models of obesity and diabetes mellitus. Nat Rev Endocrinol 2018;14:140–162.

44. Winzell MS, Ahren B. The High-Fat Diet-Fed Mouse: A Model for Studying Mechanisms and Treatment of Impaired Glucose Tolerance and Type 2 Diabetes. Diabetes 2004;53:S215–S219.

45. Turner N, Kowalski GM, Leslie SJ, Risis S, Yang C, Lee-Young RS et al. Distinct patterns of tissue-specific lipid accumulation during the induction of insulin resistance in mice by high-fat feeding. Diabetologia 2013;56:1638–48.

46. Akinmokun A, Selby PL, Ramaiya K, Alberti KGMM. The Short Insulin Tolerance Test for Determination of Insulin Sensitivity: A Comparison with the Euglycaemic Clamp. Diabet Med 1992;9:432–437.

47. Hughey CC, Wasserman DH, Lee-Young RS, Lantier L. Approach to assessing determinants of glucose homeostasis in the conscious mouse. Mamm Genome 2014;25:522–538.

48. Kim Y-B, Nikoulina SE, Ciaraldi TP, Henry RR, Kahn BB. Normal insulin-dependent activation of Akt/protein kinase B, with diminished activation of phosphoinositide 3-kinase, in muscle in type 2 diabetes. J Clin Invest 1999;104:733–741.

49. Albers PH, Bojsen-Møller KN, Dirksen C, Serup AK, Kristensen DE, Frystyk J et al. Enhanced insulin signaling in human skeletal muscle and adipose tissue following gastric bypass surgery. Am J Physiol Regul Integr Comp Physiol 2015;309:R510–24.

50. Singh R, Pervin S, Lee S-J, Kuo A, Grijalva V, David J et al. Metabolic profiling of follistatin overexpression: a novel therapeutic strategy for metabolic diseases. Diabetes, Metab Syndr Obes Targets Ther 2018;Volume 11:65–84.

51. Akpan I, Goncalves MD, Dhir R, Yin X, Pistilli EE, Bogdanovich S et al. The effects of a soluble activin type IIB receptor on obesity and insulin sensitivity. Int J Obes (Lond) 2009;33:1265–73.

52. Guo T, Jou W, Chanturiya T, Portas J, Gavrilova O, McPherron AC. Myostatin inhibition in muscle, but not adipose tissue, decreases fat mass and improves insulin sensitivity. PLoS One 2009;4:e4937.

53. McPherron AC, Lee S-J. Suppression of body fat accumulation in myostatin-deficient mice. J Clin Invest 2002;109:595–601.

54. Goncalves MD, Pistilli EE, Balduzzi A, Birnbaum MJ, Lachey J, Khurana TS et al. Akt Deficiency Attenuates Muscle Size and Function but Not the Response to ActRIIB Inhibition. PLoS One 2010;5:e12707.

55. Tao R, Wang C, Stöhr O, Qiu W, Hu Y, Miao J et al. Inactivating hepatic follistatin alleviates hyperglycemia. Nat Med 2018;24:1058–1069.

56. Patel K. Follistatin. Int J Biochem Cell Biol 1998;30:1087–1093.

57. Rae K, Hollebone K, Chetty V, Clausen D, McFarlane J. Follistatin serum concentrations during full-term labour in women – significant differences between spontaneous and induced labour. Reproduction 2007;134:705–711.

58. Vamvini MT, Aronis KN, Chamberland JP, Mantzoros CS. Energy deprivation alters in a leptin- and cortisol-independent manner circulating levels of activin A and follistatin but not myostatin in healthy males. J Clin Endocrinol Metab 2011;96:3416–23.

59. Hansen JS, Rutti S, Arous C, Clemmesen JO, Secher NH, Drescher A et al. Circulating Follistatin Is Liver-Derived and Regulated by the Glucagon-to-Insulin Ratio. J Clin Endocrinol Metab 2016;101:550–60.

60. Hansen J, Rinnov A, Krogh-Madsen R, Fischer CP, Andreasen AS, Berg RMG et al. Plasma follistatin is elevated in patients with type 2 diabetes: relationship to hyperglycemia, hyperinsulinemia, and systemic low-grade inflammation. Diabetes Metab Res Rev 2013;29:463–472.

61. Yndestad A, Haukeland JW, Dahl TB, Bjøro K, Gladhaug IP, Berge C et al. A complex role of activin A in non-alcoholic fatty liver disease. Am J Gastroenterol 2009;104:2196–205.

62. Gilson H, Schakman O, Kalista S, Lause P, Tsuchida K, Thissen J-P. Follistatin induces muscle hypertrophy through satellite cell proliferation and inhibition of both myostatin and activin. Am J Physiol Metab 2009;297:E157–E164.

63. Bodine SC, Stitt TN, Gonzalez M, Kline WO, Stover GL, Bauerlein R et al. Akt/mTOR pathway is a crucial regulator of skeletal muscle hypertrophy and can prevent muscle atrophy in vivo. Nat Cell Biol 2001;3:1014–1019.

64. Rommel C, Bodine SC, Clarke BA, Rossman R, Nunez L, Stitt TN et al. Mediation of IGF-1-induced skeletal myotube hypertrophy by PI(3)K/Akt/mTOR and PI(3)K/Akt/GSK3 pathways. Nat Cell Biol 2001;3:1009–1013.

65. Chang PY, Le Marchand-Brustel Y, Cheatham LA, Moller DE. Insulin stimulation of mitogen-activated protein kinase, p90rsk, and p70 S6 kinase in skeletal muscle of normal and insulin-resistant mice. Implications for the regulation of glycogen synthase. J Biol Chem 1995;270:29928–35.

66. Kalista S, Schakman O, Gilson H, Lause P, Demeulder B, Bertrand L et al. The type 1 insulin-like growth factor receptor (IGF-IR) pathway is mandatory for the follistatin-induced skeletal muscle hypertrophy. Endocrinology 2012;153:241–53.

67. Chevalier S, Marliss EB, Morais JA, Lamarche M, Gougeon R. Whole-body protein anabolic response is resistant to the action of insulin in obese women. Am J Clin Nutr 2005;82:355–365.

68. Castets P, Lin S, Rion N, Di Fulvio S, Romanino K, Guridi M et al. Sustained activation of mTORC1 in skeletal muscle inhibits constitutive and starvation-induced autophagy and causes a severe, late-onset myopathy. Cell Metab 2013;17:731–44.

69. Sylow L, Kleinert M, Richter EA, Jensen TE. Exercise-stimulated glucose uptake-regulation and implications for glycaemic control. Nat Rev Endocrinol 2017;13.

70. Mafakheri S, Chadt A, Al-Hasani H. Regulation of RabGAPs involved in insulin action. Biochem Soc Trans 2018;46:683–690.

71. Dokas J, Chadt A, Nolden T, Himmelbauer H, Zierath JR, Joost H-G et al. Conventional knockout of Tbc1d1 in mice impairs insulin- and AICAR-stimulated glucose uptake in skeletal muscle. Endocrinology 2013;154:3502–14.

72. Pehmøller C, Treebak JT, Birk JB, Chen S, Mackintosh C, Hardie DG et al. Genetic disruption of AMPK signaling abolishes both contraction- and insulin-stimulated TBC1D1 phosphorylation and 14-3-3 binding in mouse skeletal muscle. Am J Physiol Endocrinol Metab 2009;297:E665–75.

73. Levinger I, Jerums G, Stepto NK, Parker L, Serpiello FR, McConell GK et al. The effect of acute exercise on undercarboxylated osteocalcin and insulin sensitivity in obese men. J Bone Miner Res 2014;29:2571–2576.

74. Ikeda S, Tamura Y, Kakehi S, Takeno K, Kawaguchi M, Watanabe T et al. Exercise-induced enhancement of insulin sensitivity is associated with accumulation of M2-polarized macrophages in mouse skeletal muscle. Biochem Biophys Res Commun 2013;441:36–41.

75. Pehmøller C, Brandt N, Birk JB, Høeg LD, Sjøberg KA, Goodyear LJ et al. Exercise alleviates lipid-induced insulin resistance in human skeletal muscle-signaling interaction at the level of TBC1 domain family member 4. Diabetes 2012;61:2743–52.

76. Treebak JT, Frøsig C, Pehmøller C, Chen S, Maarbjerg SJ, Brandt N et al. Potential role of TBC1D4 in enhanced post-exercise insulin action in human skeletal muscle. Diabetologia 2009;52:891–900.

77. Mendell JR, Sahenk Z, Malik V, Gomez AM, Flanigan KM, Lowes LP et al. A phase 1/2a follistatin gene therapy trial for becker muscular dystrophy. Mol Ther 2015;23:192–201.

78. von Haehling S, Morley JE, Coats AJS, Anker SD. Ethical guidelines for publishing in the journal of cachexia, sarcopenia and muscle: update 2017. J Cachexia Sarcopenia Muscle 2017;8:1081–1083.

